# The association of child maltreatment and systemic inflammation in adulthood: A systematic review

**DOI:** 10.1101/2020.11.30.403659

**Authors:** Daniel M Kerr, James McDonald, Helen Minnis

## Abstract

**Introduction:** Child maltreatment (CM) is associated with mental and physical health disorders in adulthood. Some studies have identified elevated markers of systemic inflammation in adult survivors of CM, and inflammation may mediate the association between CM with later health problems. However, there are methodological inconsistencies in studies of the association between CM and systemic inflammation and findings are conflicting. We performed a systemic review to examine the association of CM with systemic inflammation in adults.

**Methods:** A systematic review was performed following PRISMA . Medline, Embase, Scopus and PsychInfo were searched for studies of the association of CM with blood markers of inflammation in adults. Quality was assessed using the Crowe Critical Appraisal Tool. We had intended to perform a meta-analysis but this was not possible due to variation in study design and reporting.

**Results:** Forty-six articles met criteria for inclusion in the review. The most widely reported biomarkers were C-Reactive Protein (CRP) (n=29), interleukin-6 (n=25) and Tumour Necrosis Factor-alpha (TNF-a). Four studies were prospective (all relating to CRP) and the remainder were retrospective. 85% of studies were based in Western settings. In the prospective studies, CM was associated with elevated CRP in adulthood. Results of retrospective studies were conflicting. Methodological issues relating to the construct of CM, methods of analysis, and accounting for confounding or mediating variables (particularly Body Mass Index) may contribute to the uncertainty in the field.

**Conclusions:** There is some robust evidence from prospective studies that CM is associated with elevated CRP in adulthood. We have identified significant methodological issues in the literature and have proposed measures that future researchers could employ to improve consistency across studies. Further prospective, longitudinal, research using robust and comparable measures of CM with careful consideration of confounding and mediating variables are required to bring clarity to this field.

## Introduction

Childhood maltreatment (CM) is common worldwide[1, 2]. Studies have consistently shown CM, particularly multiple and cumulative exposures, to be associated with a range of adverse physical, psychological, and social outcomes[1–6]. That this association persists after adjustment for environmental and behavioural factors suggests underlying biological mechanisms which may mediate the relationship between CM and health and social outcomes in later life[2, 7]. Understanding the biological correlates of CM will help to clarify the mechanisms linking CM with adverse outcomes, and offers the prospect of enhanced risk stratification of young people who have been subject to maltreatment and may identify new treatment targets to break the link between childhood experiences and adverse physical and mental health outcomes in adulthood[2].

Low-grade systemic inflammation is generally defined as 2-3 fold elevations in inflammatory markers like C-reactive protein (CRP), Interleukin-6 (Il-6) and tumour necrosis factor alpha (TNF-a)[8]. This represents a chronic low-level activation of the immune system (likely representing excessive sensitivity to inflammatory stimuli and deficiencies of the anti-inflammatory pathways which would normally terminate such responses) and is distinguished from high-grade inflammatory states with markedly elevated inflammatory markers such as occurs in acute infections, severe illnesses, and auto-inflammatory diseases. Low-grade, systemic inflammation has been identified in adult survivors of CM[9].

### Inflammation and physical health disorders

Low-grade inflammation has been associated with a range of physical health conditions such as cardiovascular disease and diabetes[10, 11]. Notably, a large body of work has associated low-grade elevations in CRP with cardiovascular events - however subsequent work has questioned the direction of causality in this relationship[12]. Other inflammatory markers have been associated with cardiovascular disease, particularly Interleukin-6[10]. A large international study using Mendelian randomisation techniques has supported a causal relationship between elevated levels of Il-6 and cardiac disease[10]. Further supporting evidence for the role of low-grade inflammation in cardiovascular disease is provided by the recent CANTO trial of the specific Il-1b antagonist Canakinumab which was shown to reduce rates of myocardial infarction, stroke and death in patients treated following an MI with elevated CRP[13].

### Inflammation and mental health disorders

Low-grade inflammation is also associated with a range mental health disorders. A wide body of work has associated major depressive disorder with low-grade elevations in inflammatory markers like CRP, Il-6, and TNF-a[14, 15]. The neurobiological effects of peripheral cytokines may mediate the relationship between external stressors and depression[14]. Low-grade inflammation is also associated with conditions like post-traumatic stress disorder (PTSD), schizophrenia, and bipolar affective disorder[8, 16]. A recent Mendelian randomisation analysis has suggested a causal relationship between CRP and both schizophrenia and bipolar affective disorder[16].

### Inflammation and child maltreatment

An emerging body of evidence, therefore, has shown that low-grade systemic inflammation is associated with physical and mental health disorders and that this relationship is at least partially causal. Although it is known that CM is associated with low-grade, systemic inflammation [9], previous reviews have identified significant heterogeneity in the literature particularly in relation to the definition and ascertainment of CM [17, 18]. Studies have offered varying definitions of CM ranging from narrowly focused childhood physical or sexual abuse, to more broadly defined Adverse Childhood Experience (ACEs)[17, 18]. These differing patterns of CM will likely have different effects on development, contributing to the heterogeneity in the literature. Furthermore, research in this area has highlighted the role of potential mediators between CM and inflammation (particularly body mass index (BMI)) and raise questions about the causality of this relationship that were not fully addressed in previous reviews[19, 20]. In this light, this current review aimed to examine the association between CM and systemic inflammation in adulthood, with particular consideration of the causality of this relationship.

## Methods

We performed a systematic review of the association between CM and low-grade inflammation. Preferred Reporting Items for Systematic Reviews and Meta-Analyses (PRISMA) guidelines were followed[21]. The search and synthesis plan were pre-specified in a protocol registered with PROSPERO (CRD42020187027).

### Research questions

Our primary research question was: “Is CM associated with elevated markers of systemic inflammation adulthood?”. We also prespecified secondary questions: “Are differences in later life inflammation associated with specific sub-types, timings, or durations of abuse?” and “What mediates any association between CM and inflammation?”

### Inclusion criteria

We included non-randomised observational studies. Our study population was human adults (>18 years of age). Participants could be drawn from healthy samples or clinical (mental or physical health disorder) samples, however we excluded studies of participants with pro-inflammatory physical health conditions (in particular autoimmune disease and cancer). Participants could be drawn from community or hospital-based samples.

Our exposure was CM (defined as physical abuse, sexual abuse, emotional abuse, physical neglect and/or emotional neglect occurring at least once before the age of 18). We did not place any restrictions on how CM was recorded. Studies could use retrospective or prospective ascertainment; and could record CM using validated scales, or specific measures developed for their study if this was described. CM could be reported as an overall construct or broken down into sub-types of abuse and neglect. Studies could compare between a CM exposed group and a control group or utilise a continuous measure of CM in a sample.

Our outcome was blood levels of inflammatory markers. Any marker of the inflammatory response measured in the blood was eligible for inclusion. We excluded studies which reported on stimulated responses (e.g. to stress testing or biological stimulation), studies reporting exclusively on gene expression, *in vitro* production of inflammatory mediators, and studies exclusively measuring inflammation in the central nervous system (e.g. CSF).

### Search strategy

We searched MedLine, Embase, PsychInfo, and Scopus. Our first search term aimed to capture CM. This included MeSH terms “child abuse”, “child abuse, sexual”, “adult survivors of child abuse”, “physical abuse”, “child, abandoned”, “adolescent, institutionalized”, “adult survivors of child adverse events” and “adverse childhood experiences”, supplemented by title and abstract searches for related terms. The second term sought to identify broadly defined inflammatory dysfunction. This included MeSH terms “inflammation”, “C-reactive protein”, “acute phase proteins”, “tumour necrosis factor alpha”, “interleukins”, “cytokines”, “immune system”, “fibrinogen”, “leukocytes”, and “lymphocytes”. This was supplemented by title and abstract searches for related terms.

These terms were combined using the Boolian operator “AND”, and duplicates were removed.

The search was adapted to utilise relevant keywords in the other databases used and full information is available in S1 file.

This search was supplemented by manual checking of reference lists of retrieved articles and checking the reference lists of previous reviews in this area.

## Methods of review

Records were initially screened against inclusion criteria by 1 reviewer, and a 2^nd^ reviewer independently reviewed a sub-sample of 25% of titles. All included articles were then reviewed by a 2^nd^ reviewer to confirm that they met inclusion criteria. Disagreements were resolved through conference with a 3^rd^ author.

Risk of bias assessment was performed at study level using the Crowe Critical Appraisal Tool v1.4 (https://conchra.com.au/wp-content/uploads/2015/12/CCAT-form-v1.4.pdf). This is a tool for assessment of risk of bias in non-randomised studies. Key components of risk of bias assessment include sampling, ascertainment of exposure, measurement of outcome, and statistical analysis including adjustment for relevant confounding variables. The CCAT assigns a total score from 0-40. According to the tools guidelines studies can be categorised as low quality (<20), moderate quality (20-29), and high quality (30+). All papers were rated by 1 reviewer, and the 2^nd^ reviewer independently quality rated a sub-sample of 25% of papers. Again, disagreements were resolved through conference with a 3^rd^ author.

Data extraction was performed using a pre-specified form. Data extraction was performed by 1 reviewer, with a 2^nd^ reviewer independently performing data extraction for a sample of 25% of included papers.

### Synthesis

We had intended to perform a meta-analysis of the most widely reported inflammatory markers as specified in the protocol. A more detailed review of included articles showed that this was not feasible due to differences in the construct of CM being utilised, incommensurable methods of analysis and inconsistent accounting for covariates. This is discussed further below. These challenges led us to conclude that a meta-analysis would be of questionable validity. We have instead presented our findings in a narrative format with focus on the most widely reported inflammatory markers, and methodological factors.

## Results

The PRISMA flow chart is shown in figure 1. Details of reasons for exclusion of articles are shown in S1 table. A total of 46 papers were included in this review. All papers were rated as moderate to high quality (CCAT scores ranged from 23-38, median- 32). 30 papers reported on multiple biomarkers. The frequency with which biomarkers were reported is shown in table 1. The most widely reported biomarkers were CRP (n=29), interleukin-6 (n=25), and TNF-a (n= 16). Details of these are discussed below. Details of other biomarkers reported are shown in S2 table.

**Figure 1.**
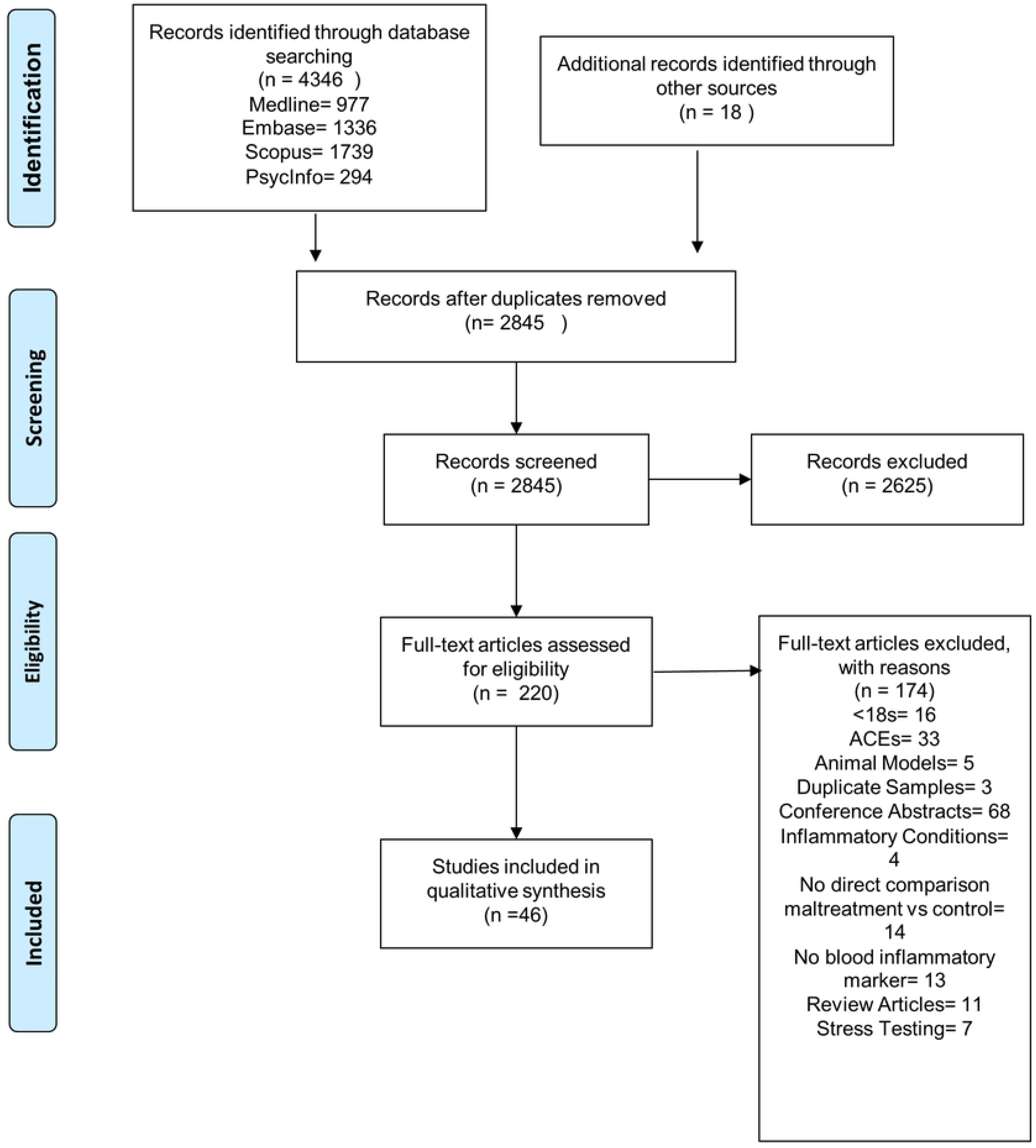
PRISMA Flow Diagram

**Table 1.** Frequency with which biomarkers are reported in included studies

| Biomarker | Number of studies |
| --- | --- |
| C-reactive protein (CRP) | 29 |
| Interleukin 6 (IL-6) | 25 |
| Tumour necrosis factor alpha (TNF-a) | 16 |
| Interleukin 1b (IL-1b) | 8 |
| Interleukin 10 (IL-10) | 6 |
| Fibrinogen | 5 |
| Soluble tumour necrosis factor receptor 1 (sTNFR1) | 3 |
| Soluble tumour necrosis factor receptor 2 (sTNFR2) | 2 |
| Adiponectin | 2 |
| Brain derived neurotrophic factor (BDNF) | 2 |
| Interferon gamma (IFN-γ) | 2 |
| Interleukin-4 (IL-4) | 2 |
| Resistin | 2 |
| Transforming growth factor beta (TGFB) | 2 |
| White cell count (WCC) | 2 |
| Interleukin 2 (IL-2) | 1 |
| Interleukin 8 (IL-8) | 1 |
| Interleukin 12 (IL-12) | 1 |
| Interleukin 1 receptor antagonist (IL-1RA) | 2 |
| Soluble interleukin 6 receptor (sIL-6R) | 1 |
| Agouti related protein (AgRP) | 1 |
| Basic fibroblast growth factor (bFGF) | 1 |
| Betacellulin (BTC) | 1 |
| Cell-mediated immunity (EBV titre) | 1 |
| Chemokine Ligand-2 (CL-2) | 1 |
| D-Dimer | 1 |
| E-Selectin | 2 |
| Glucocorticoid induced tumour necrosis factor ligand (GITR-L) | 1 |
| Glycoprotein 130 | 1 |
| Insulin-like growth factor-binding protein 2 (IGFBP2) | 1 |
| Interferon induced T-cell alpha chemoattractant (I-TAC) | 1 |
| Monocyte chemoattractant protein 1 (MCP-1) | 1 |
| Mucosae-associated epithelial chemokine (MEC) | 1 |
| Neurotrophin-4 | 1 |
| Platelet derived growth factor (PDGF) | 1 |
| Salivary alpha amylase (sAA) | 1 |
| Stem cell factor (SCF) | 1 |
| Soluble intracellular adhesion molecule 2 (sICAM2) | 1 |
| Soluble immunoglobulin A (sIgA) | 1 |
| Thymus expressed chemokine (TECK) | 1 |
| Tumour necrosis factor related apoptosis inducing ligand-receptor 4 (TRAIL-R4) | 1 |
| Vascular endothelial growth factor (VEGF) | 1 |
| Von Willebrand Factor (vWF) | 1 |

## Methodological features

Over 90% (n= 42) of the 46 included studies recorded CM exposure retrospectively, with the Childhood Trauma Questionnaire (CTQ) being the most widely reported scale (n=29). CTQ is a 28-item self-report measure of childhood trauma, which can be considered as a total score, or as subscales representing physical abuse, physical neglect, emotional abuse, emotional neglect, and sexual abuse[22]. Of papers utilising the CTQ, 13 reported only on the total score, 8 reported on subscales only, and 8 reported on both. 13 papers reported on the CTQ as a dichotomous variable (using a recognised cut-off point to define high versus low scores) and 16 analysed CTQ as a continuous variable. The remaining studies utilised their own measures of CM based on their dataset (n=9) or other standardised scales (n=8). After CTQ the most widely used standardised scale was the Early Trauma Inventory (ETI)[23]. This is a 56-item self-report scale which generates 5 variables- total number of traumas, physical trauma, emotional trauma, sexual trauma, and general traumas. General trauma includes a range of adverse exposures including parental separation, bereavement, natural disaster, and political violence. Most studies did not specify the timing or duration of CM in their analysis.

There was significant variation in statistical techniques used and where multivariate analysis was performed there was inconsistency as to which covariates were included. Eighty-five percent (n=39) of the studies were conducted in Western settings (North America, Europe, Australasia), with the remainder taking place in Brazil (n=4), China (n=2), and Japan (n=1). Ten studies were restricted to female participants, and one was restricted to males.

### C-reactive protein

The association between CM and CRP was reported in 29 papers (full details in table 1a and 1b). Nine reported on clinical samples and 20 on non-clinical samples.

**Table 1a:**
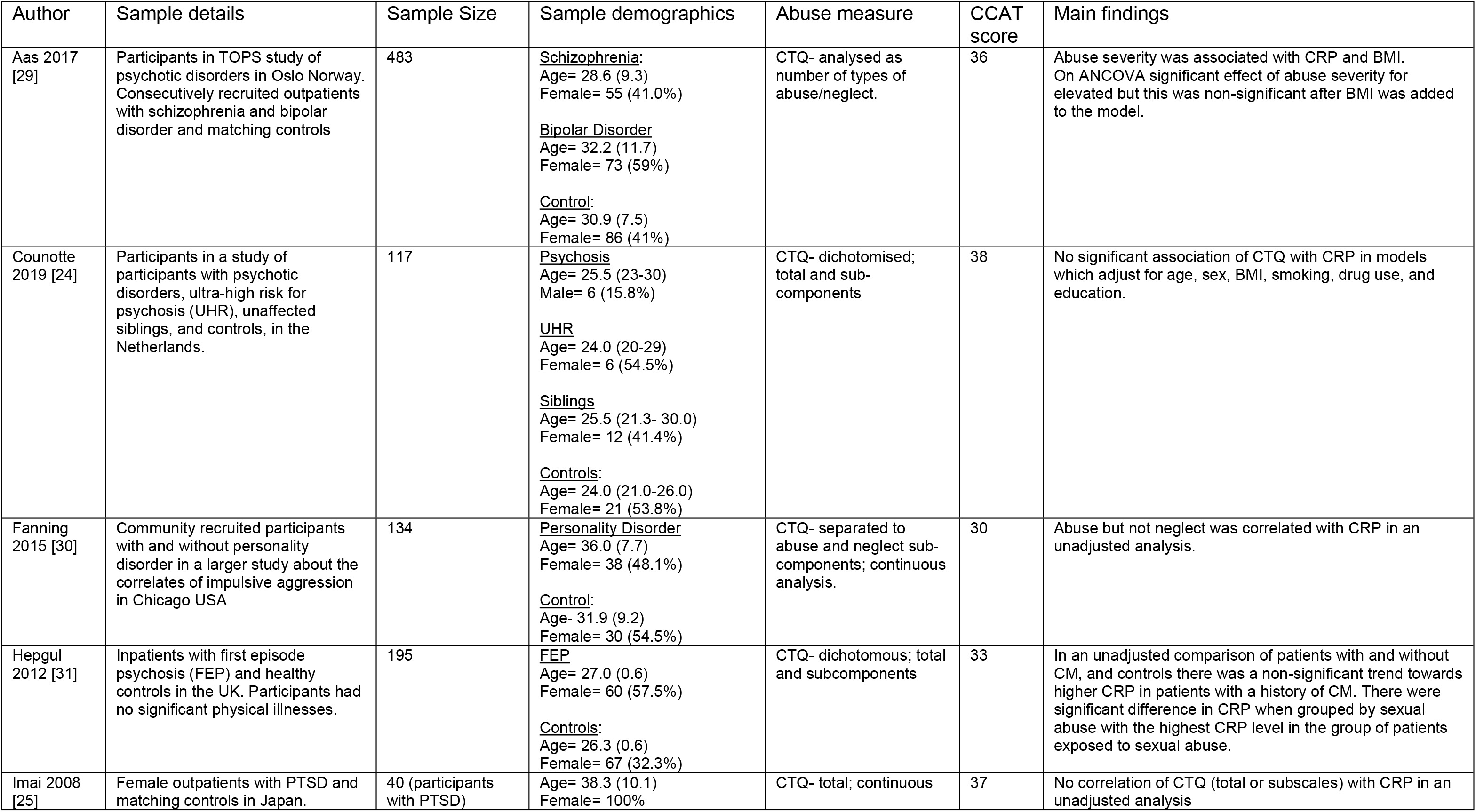
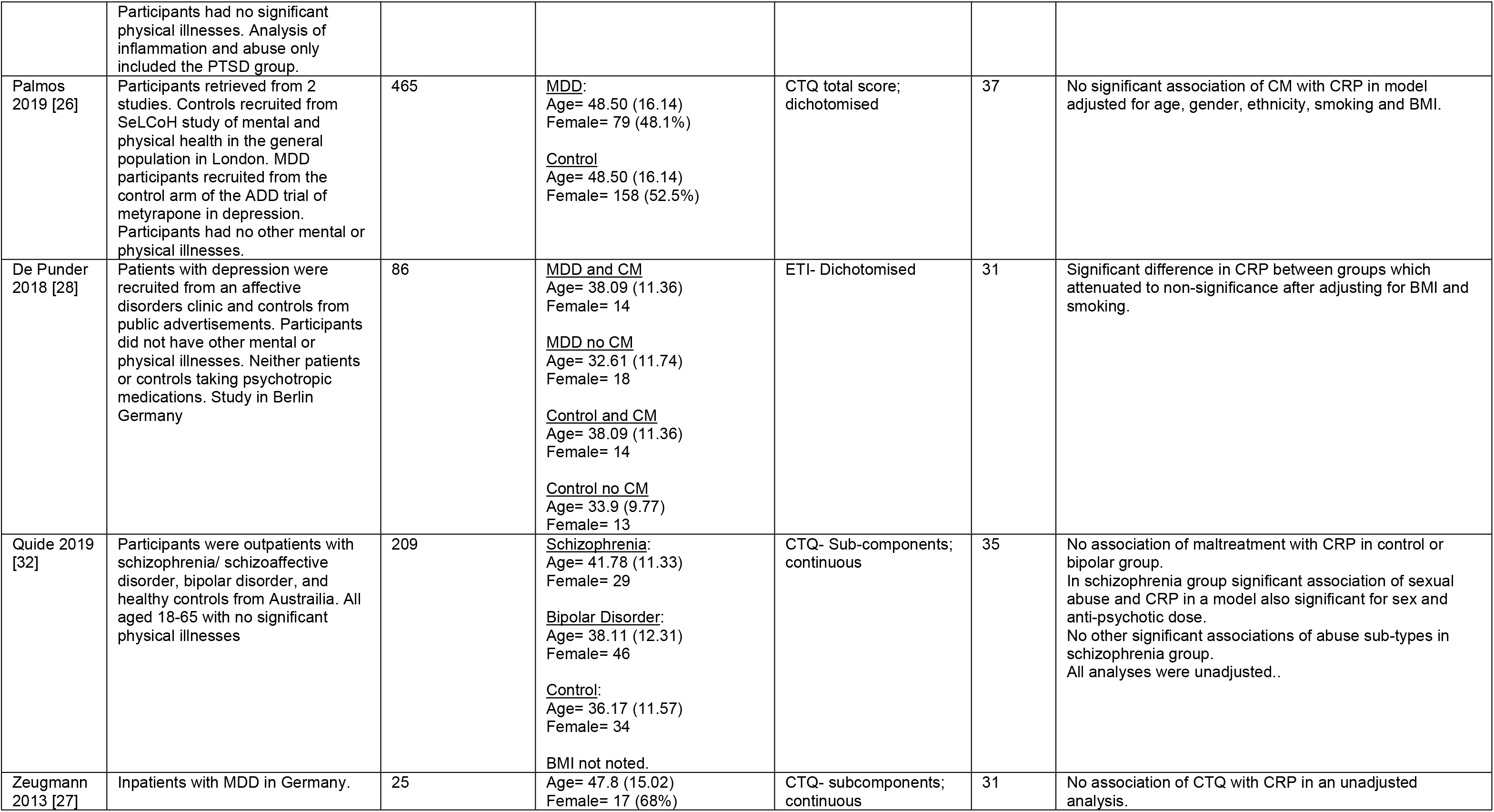
Papers reporting CRP: Clinical populations

**Table 1b.**
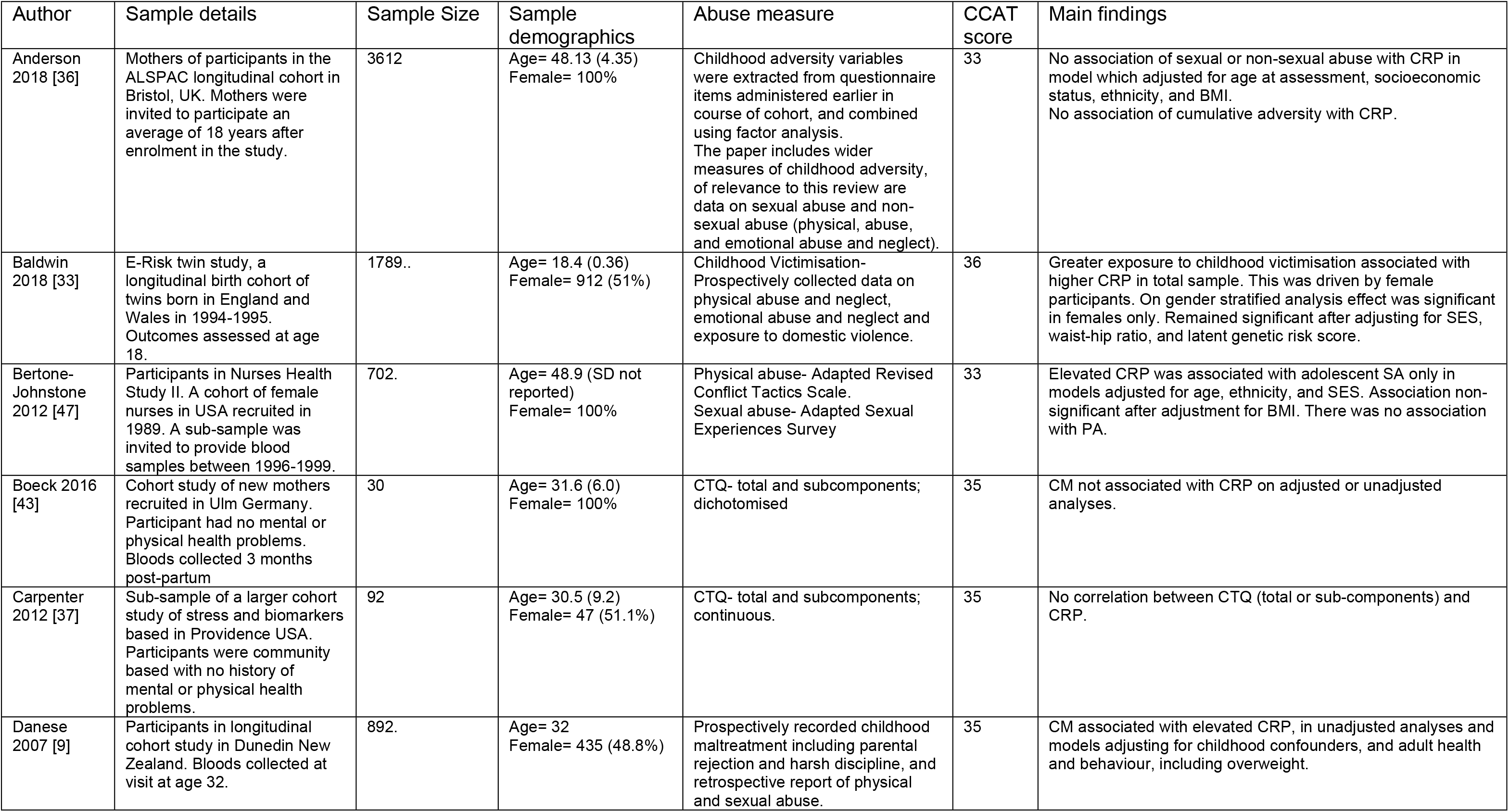
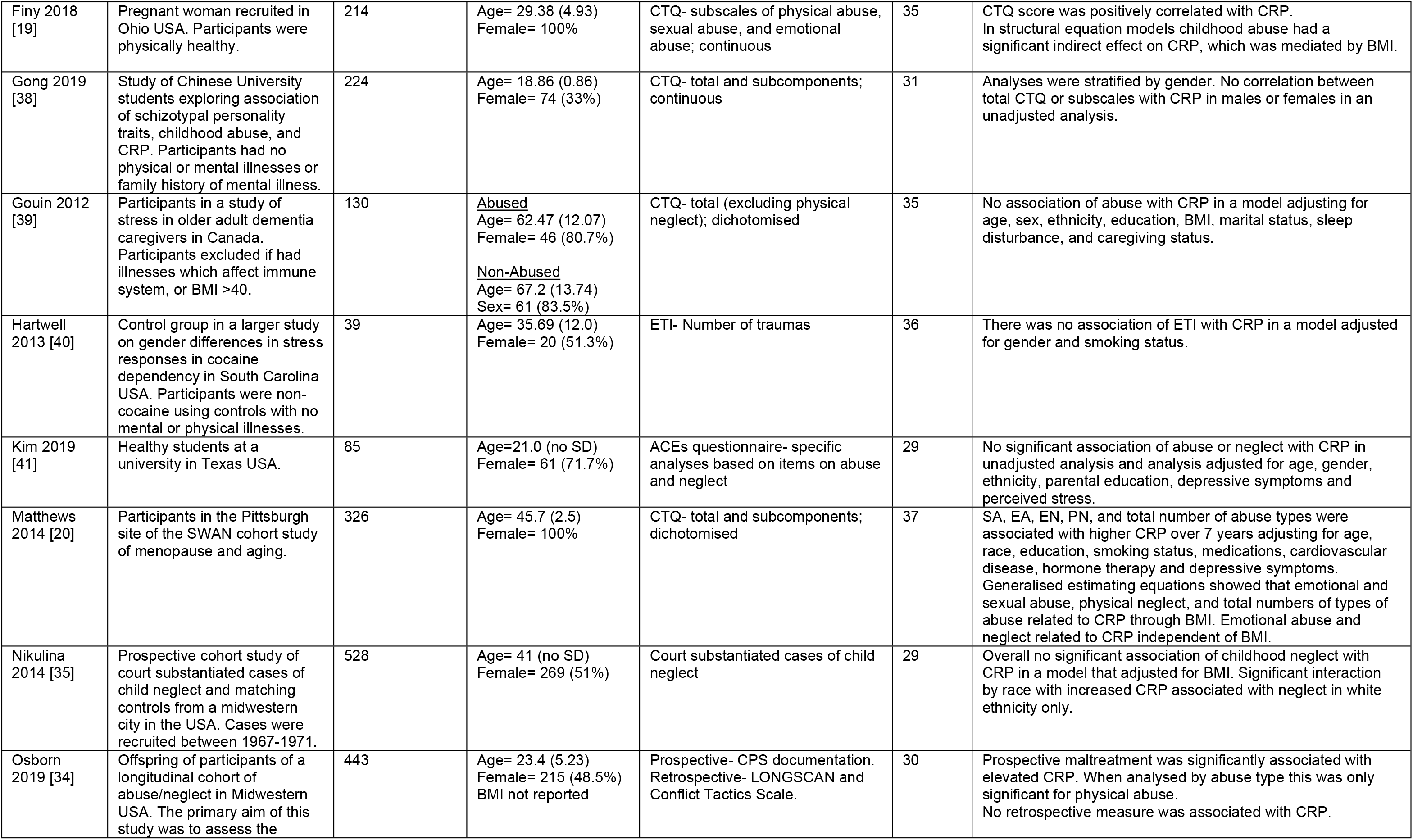
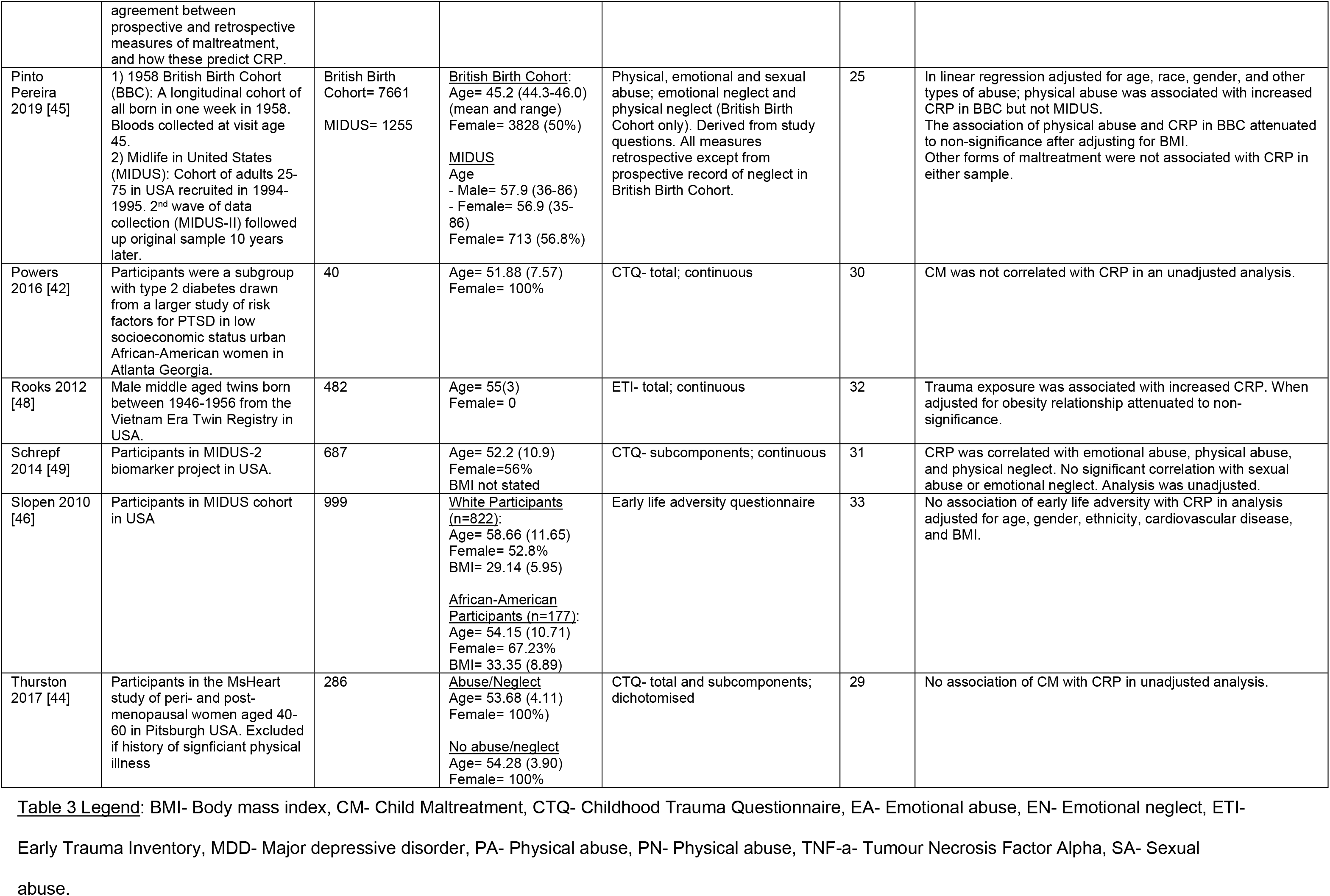
Papers reporting CRP: Non-clinical populations

All papers reporting clinical samples were retrospective and all but one used CTQ to record CM. Four studies did not find any association between CM and CRP[24–27]. A further two studies found initially significant associations between CM measures and CRP, which attenuated to non-significance on adjustment for BMI[28, 29]. Significant associations between CM with CRP were reported in three studies. In a study comparing 79 participants with personality disorder and 55 healthy controls Fanning et al demonstrated a significant association between abuse and CRP, as measured retrospectively using the CTQ, but not neglect, in a bivariate correlation which did not adjust for covariates[30]. In a sample of 96 participants with first episode psychosis and 99 healthy controls, Hepgul *et al* found a trend towards elevated CRP in patients who had experienced CM, but this only reached statistical significance when participants were grouped by exposure to sexual abuse and, again, did not adjust for covariates[31]. In a study of 209 participants with schizophrenia/schizoaffective disorder, bipolar affective disorder and healthy controls, Quide et al demonstrated a significant association between sexual abuse and elevated CRP in schizophrenia patients only[32]. No other associations of CRP with other abuse sub-types or in different clinical groups was demonstrated. The analysis adjusted for age, gender, disease severity, and medication use, but did not adjust for BMI.

Of 20 studies examining the association between CM and CRP in non-clinical samples, 16 were retrospective and four prospective. The four prospective studies found significant associations between CM and elevated CRP. In a prospective twin study, Baldwin et al reported an association between CM and elevated CRP at age 18[33]. On stratified analysis this effect was significant in females only and remained significant after adjusting for covariates including waist-hip ratio. Osborn et al reported on the association between retrospective and prospective measures of CM with CRP[34]. They found that CRP was associated with prospective measures of abuse only. Of note their analysis adjusted for age, sex, ethnicity, parental occupation, heavy drinking, smoking, and depression but did not adjust for BMI. Danese et al reported on a large prospective cohort in New Zealand that measured CM using a combination of prospective and retrospective reports[9]. They demonstrated a significant association between CM and CRP which remained significant in extensively adjusted models, including adjustments for adult health behaviours and obesity. They estimated that 10% of low-grade inflammation as measured by CRP may be independently attributable to CM. Nikulina et al report a US cohort exposed to court substantiated neglect and controls matched for age, sex, ethnicity, and socioeconomic status. In a model adjusting for BMI there was no significant association between neglect and CRP in the total sample, however the study did identify a significant interaction with race (authors’ terminology)- wherein neglect was associated with elevated CRP in white participants only[35]. Their analysis considered family poverty as a covariate in this analysis, but it did not reach significance threshold for inclusion in the final model.

Of the 16 retrospective studies in non-clinical samples, nine studies found no significant association between child maltreatment and CRP[36–44]. A further 4 found initially significant associations which attenuated to non-significance on adjustment for BMI[45–48]. Finy et al report a study of 214 pregnant women (of whom 51.4% were overweight or obese) in which they reported small but statistically significant associations of CTQ score with CRP[19]. Through structural equation modelling, this association was found to be indirect and mediated by elevated BMI. Similarly, Matthews et al in a middle-aged female sample, found that sexual abuse, physical neglect and total number of abuse types were associated with elevated CRP in a relationship mediated by elevated BMI. They also found that emotional abuse and neglect were independently associated with elevated CRP[20]. Schrepf and colleagues, in a study of 687 participants in the MIDUS-II biomarker project found that CM was significantly associated with elevated CRP in adulthood in a relationship that was mediated by elevated BMI. They found that this relationship was mediated by a latent distress measure which was associated with using food as a coping mechanism. The association between BMI and CRP was stronger in females than males[49].

To summarise an association between CM and later elevation of CRP was demonstrated in four prospective studies in non-clinical samples, which adjusted for relevant covariates (however one did not adjust for BMI). Most retrospective studies (13/16) found either no association of CRP with BMI or an association which attenuated to non-significance on adjustment for BMI or other obesity measures. Three studies found a significant association of CM with CRP, all of which found that this to be mediated by elevated BMI.

### Interleukin-6

The association between CM and Il-6 was reported in 25 papers, 15 of which were based on clinical samples. Details of included papers are shown in tables 2a and 2b. All papers utilised retrospective measures of CM. Nineteen papers used CTQ to measure CM.

**Table 2a.**
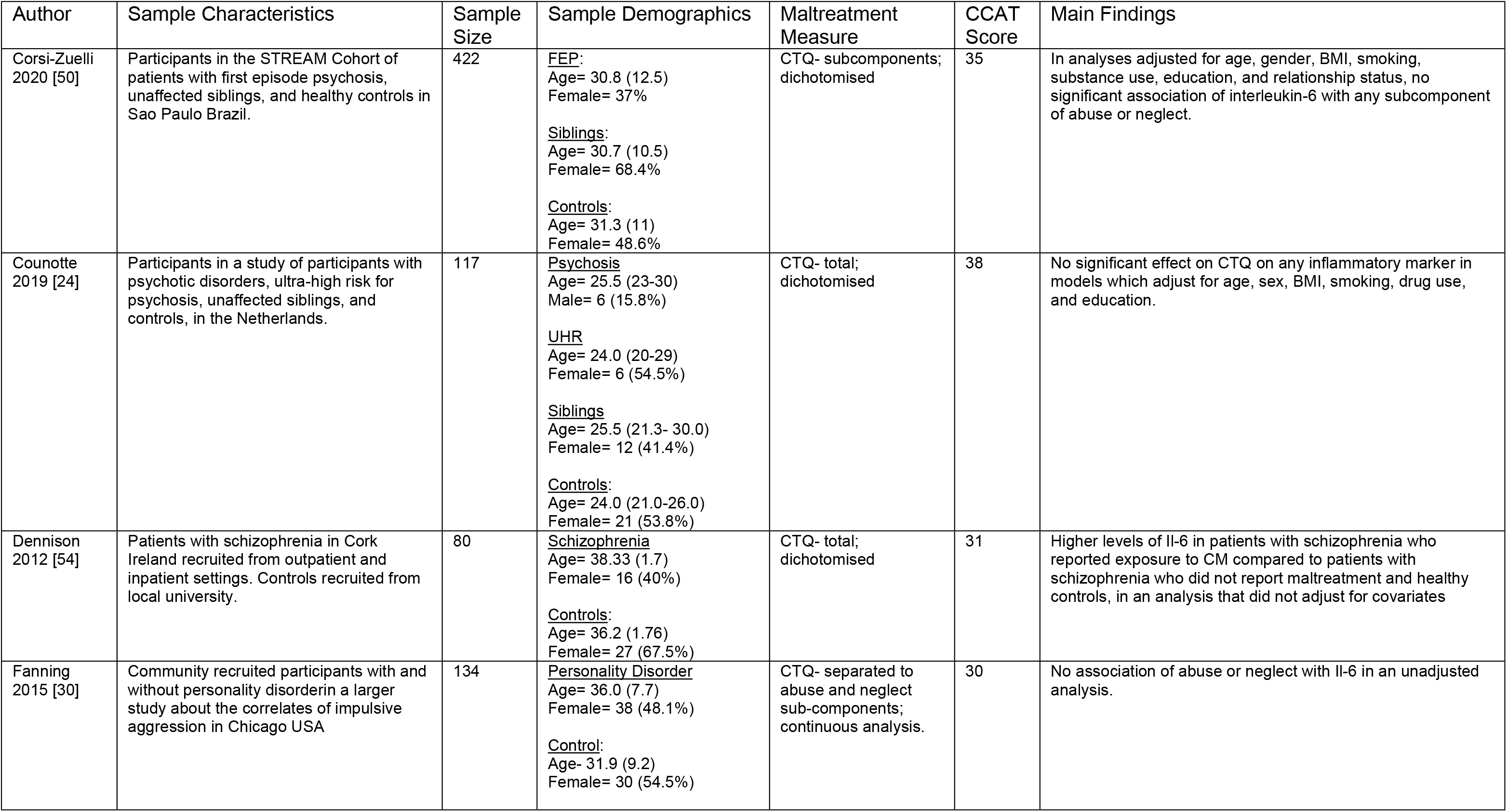
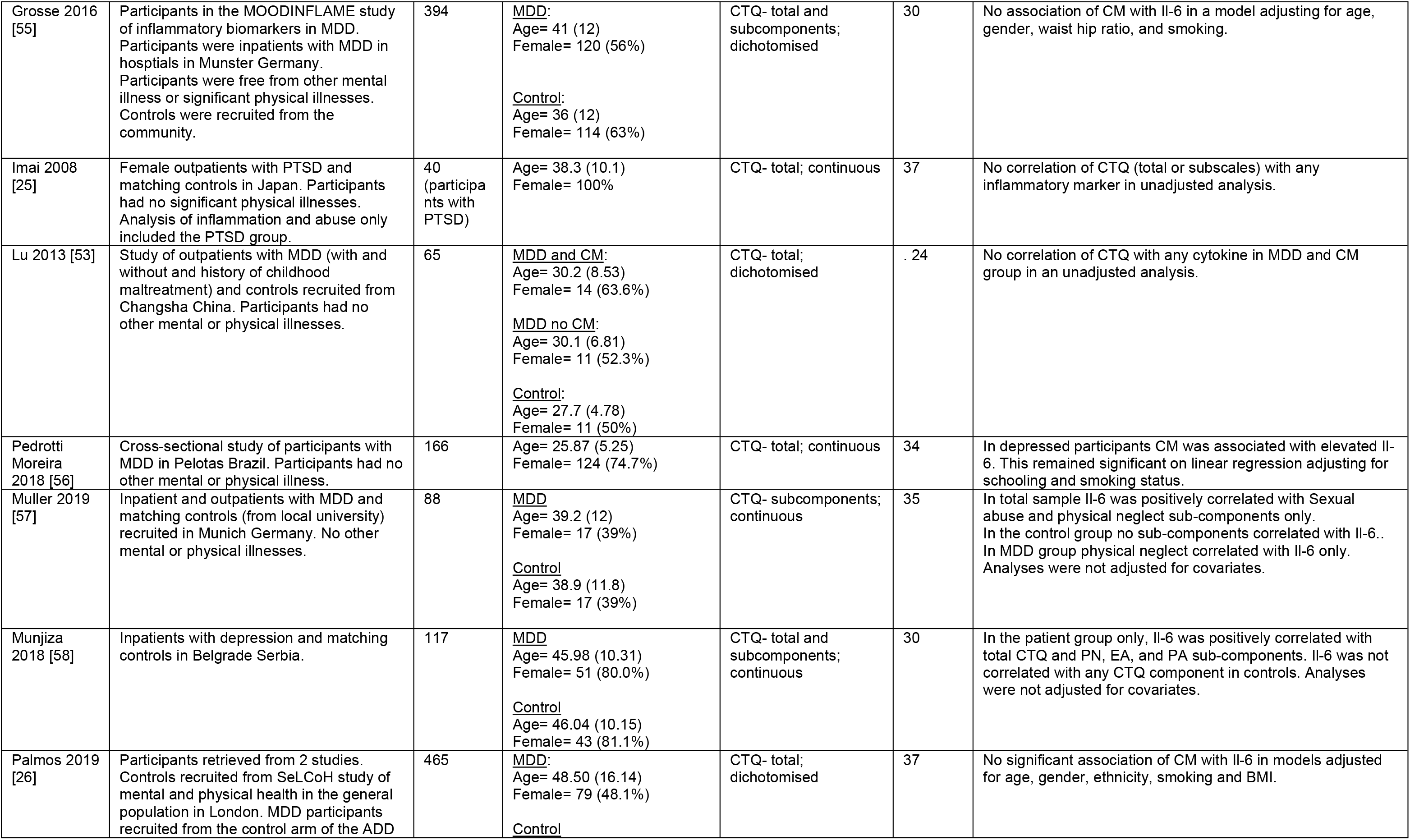
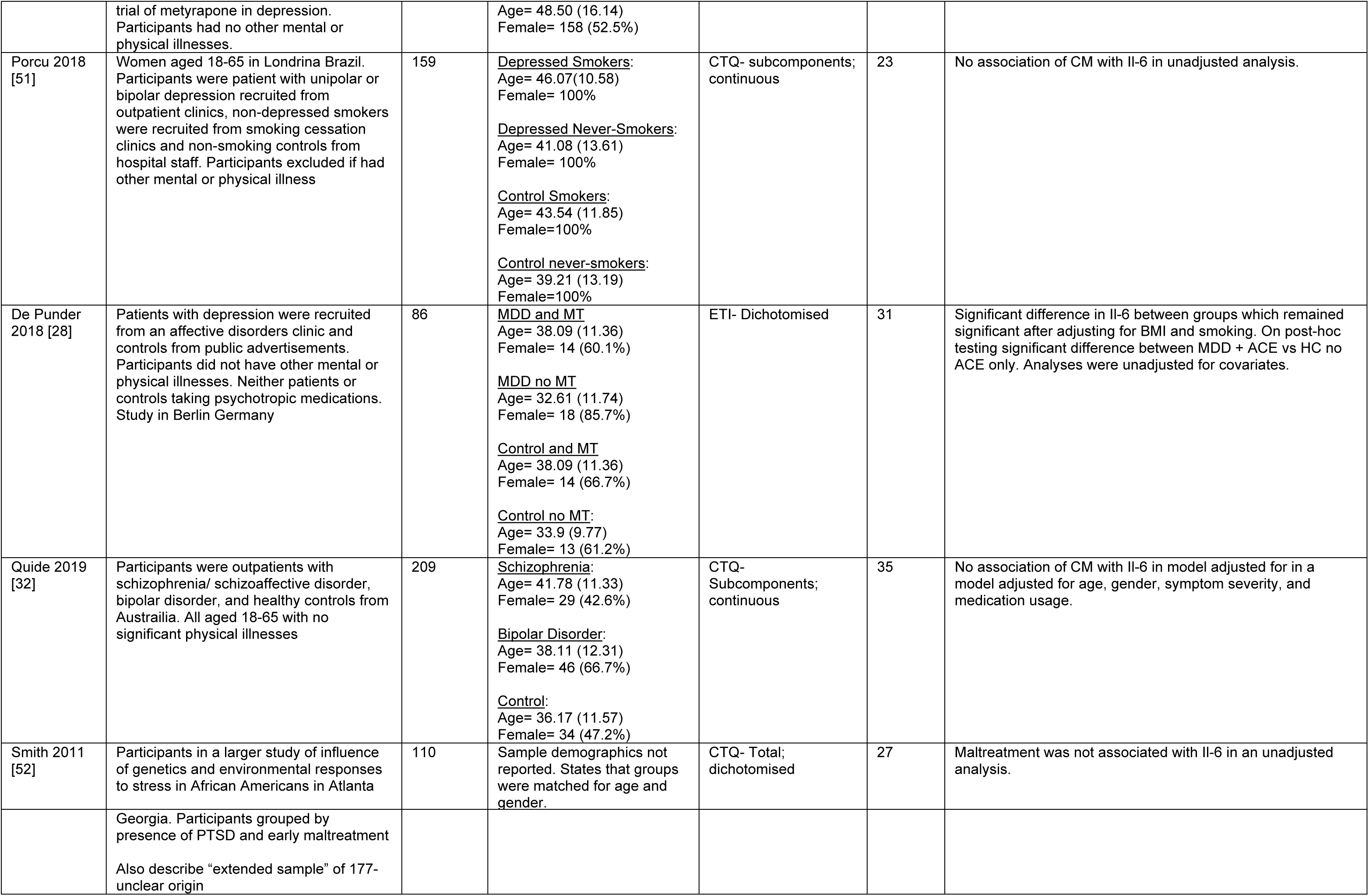
Studies reporting interleukin-6: Clinical populations

**Table 2b.**
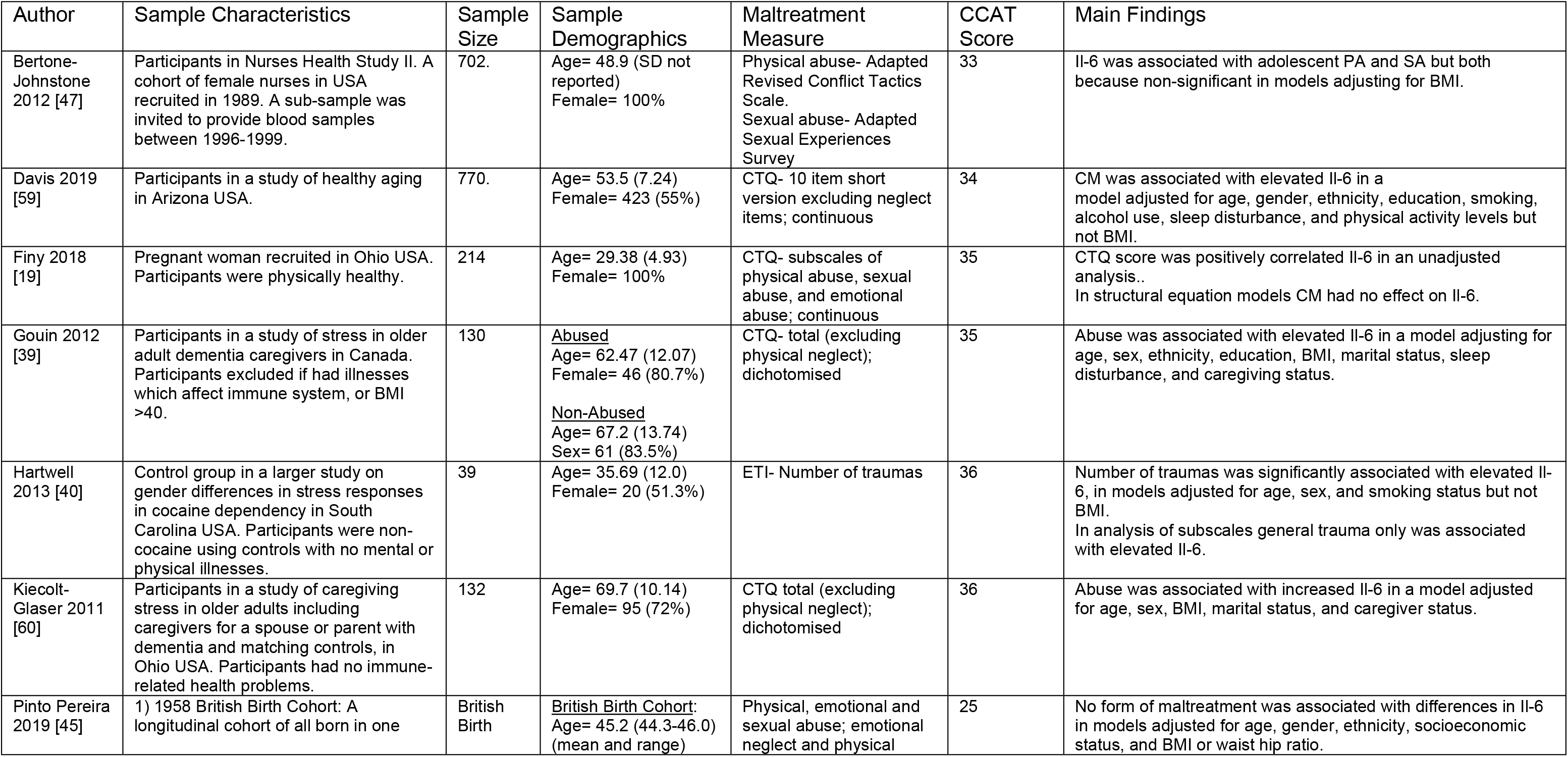
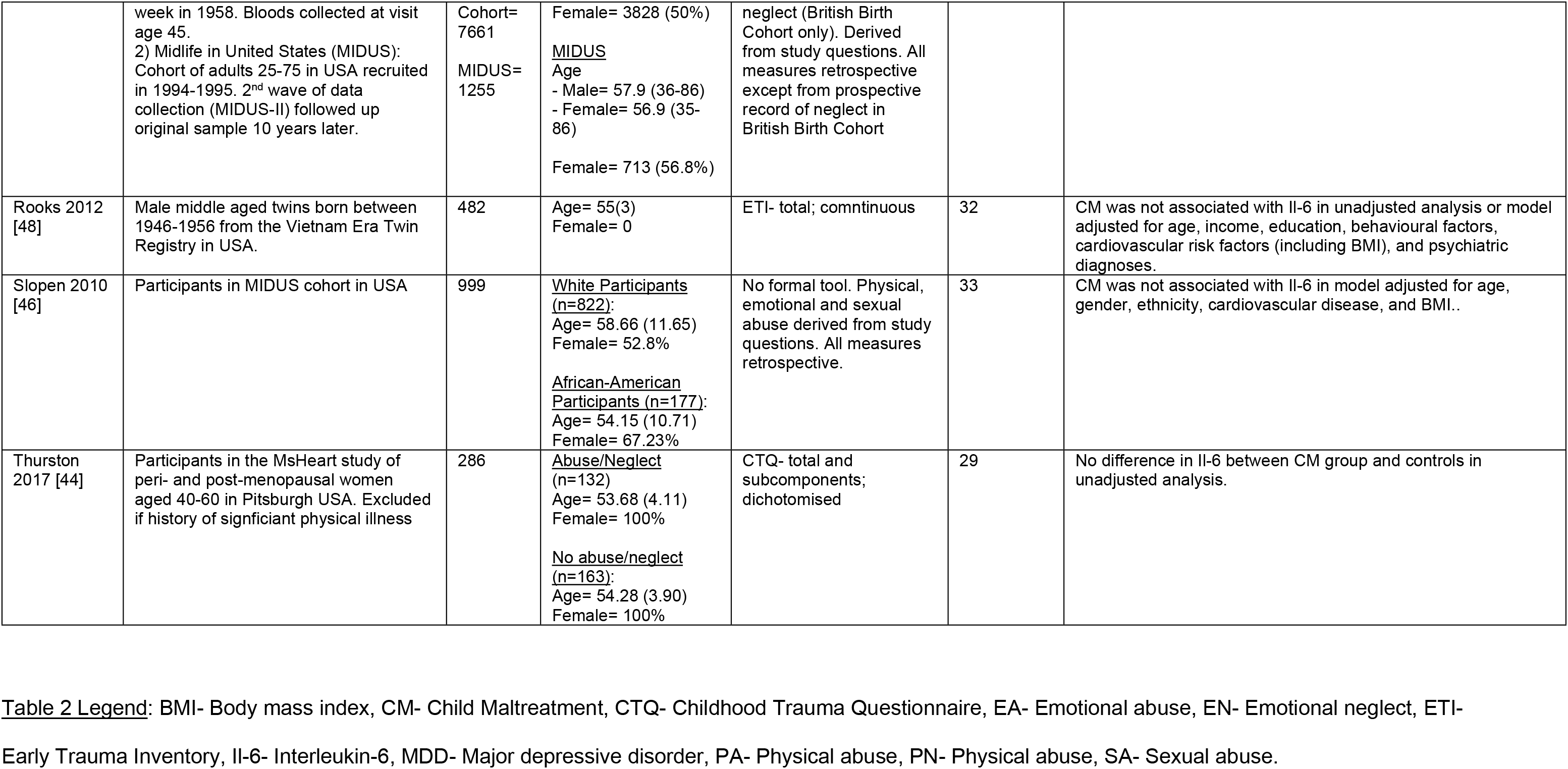
Samples reporting interleukin-6: Non-clinical populations

Of the clinical samples, nine papers did not identify a significant association between CM and Il-6[24–26, 30, 32, 50–53]. Dennison et al reported higher levels of Il-6 in patients with schizophrenia who reported exposure to CM compared to patients with schizophrenia who did not report maltreatment and healthy controls, in an analysis that did not adjust for covariates[54]. Grosse et al reported on 394 patients and controls in the MOODINFLAME study of inflammatory markers in Major Depressive Disorder (MDD)[55]. There was no association between CM and Il-6 in the total sample, nor in the MDD or control groups. In an analysis limited to MDD patients exposed to CM, sexual abuse was associated with elevated Il-6 in an analysis that adjusted for age, gender, smoking, and waist-hip ratio. Pedrotti Moreira et al reported on a cross-sectional study of MDD and healthy control participants in Brazil[56]. They identified a significant association between CM and higher Il-6 in participants with MDD only, in an analysis which adjusted for education and smoking status but not BMI. Muller et al examined the correlation between CTQ scores and inflammatory markers in a sample consisting of patients with MDD and healthy controls[57]. They found a small but significant correlation between sexual abuse and physical neglect with Il-6 in an unadjusted analysis. Munjiza reported that Il-6 was positively correlated with total CTQ, physical neglect, emotional abuse, and physical abuse in an unadjusted analysis limited to participants with MDD only[58]. De Punder et al reported on a sample of patients with MDD and healthy controls[28]. They grouped participants by presence of MDD and exposure to CM and identified a significant between group difference in an analysis which adjusted for BMI and smoking. On post-hoc testing the only significant difference was between the MDD and CM group vs healthy control and no CM, thus this analysis does not clearly distinguish the effects of MDD from CM.

Ten studies reported on non-clinical samples. Three of these found no association between CM and Il-6[45, 48]; a further three studies found associations of CM with elevated Il-6 which attenuated to non-significance after adjustment for BMI[19, 44, 46, 47]. Davis et al, in a study of healthy middle aged adults in the USA, found that CM was significantly associated with elevated Il-6 in a model that adjusted for age, gender, ethnicity, and health behaviours, but not BMI[59]. Gouin et al, in a study of care-giver stress in older adults found a significant association between CM and Il-6 in a model which adjusted for age, sex, ethnicity, education, BMI and social factors[39]. Hartwell *et al* reported a significant association between the number of traumas as measured by the ETI and elevated Il-6 in a model which adjusted for age, sex, and smoking status but not BMI[40]. In another study of care-giver stress in older adults, Kiecolt-Glaser identified a significant association between CM and elevated Il-6 in a model adjusted for age, sex, BMI and social factors[60].

In summary most studies did not find a significant association between CM and elevated Il-6. Studies reporting positive findings tended not to adjust for BMI and in some papers positive associations were limited to sub-groups. Notably, two studies that adjusted appropriately for covariates found significant associations between CM and elevated Il-6 in older adults.

### Tumour necrosis factor-alpha

The association between CM with TNF-a was reported in 16 papers, 13 of which were in clinical samples. All studies were retrospective and 14 used the CTQ to measure CM. Details of included papers are shown in tables 3a and 3b.

**Table 3a.**
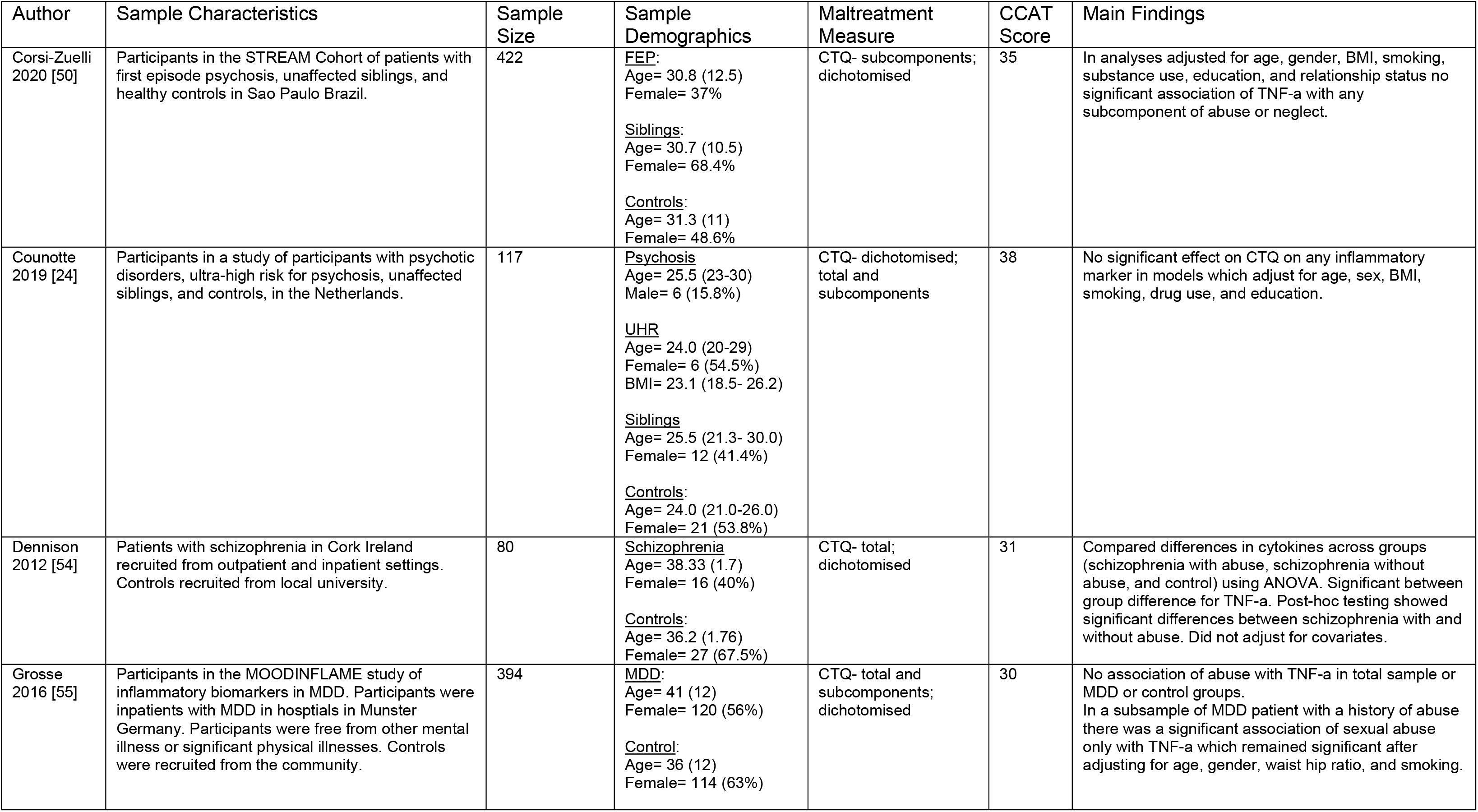
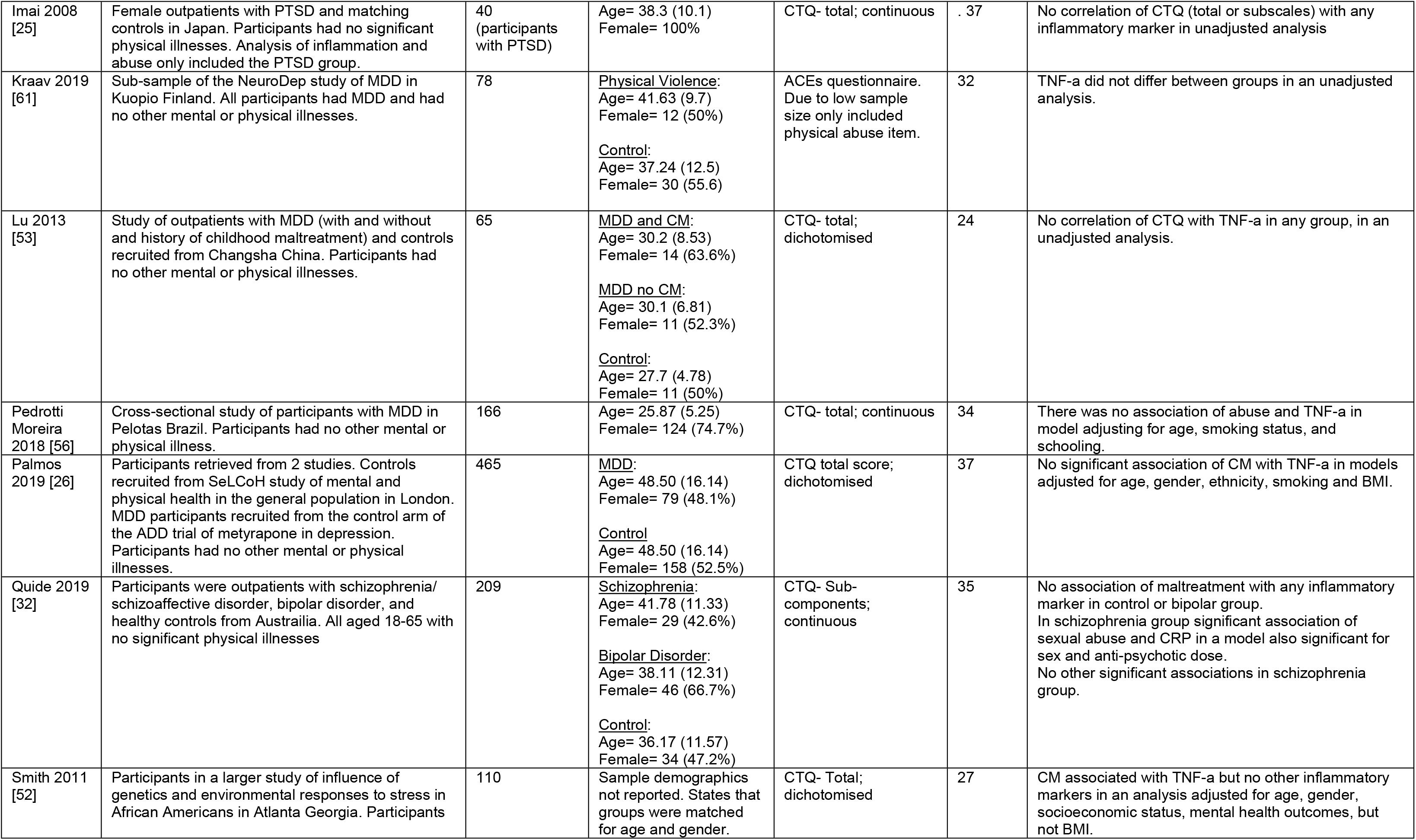
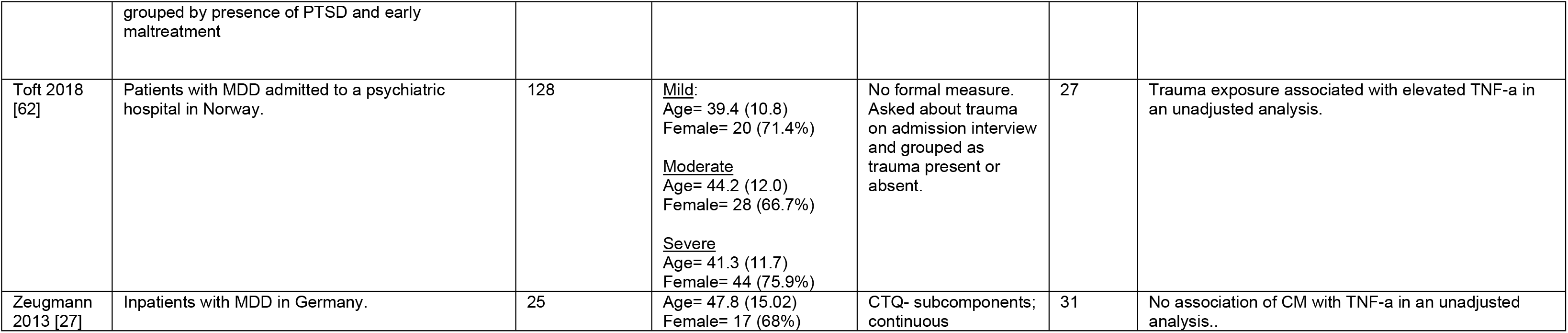
Studies reporting TNF-a: Clinical populations

**Table 3b.**
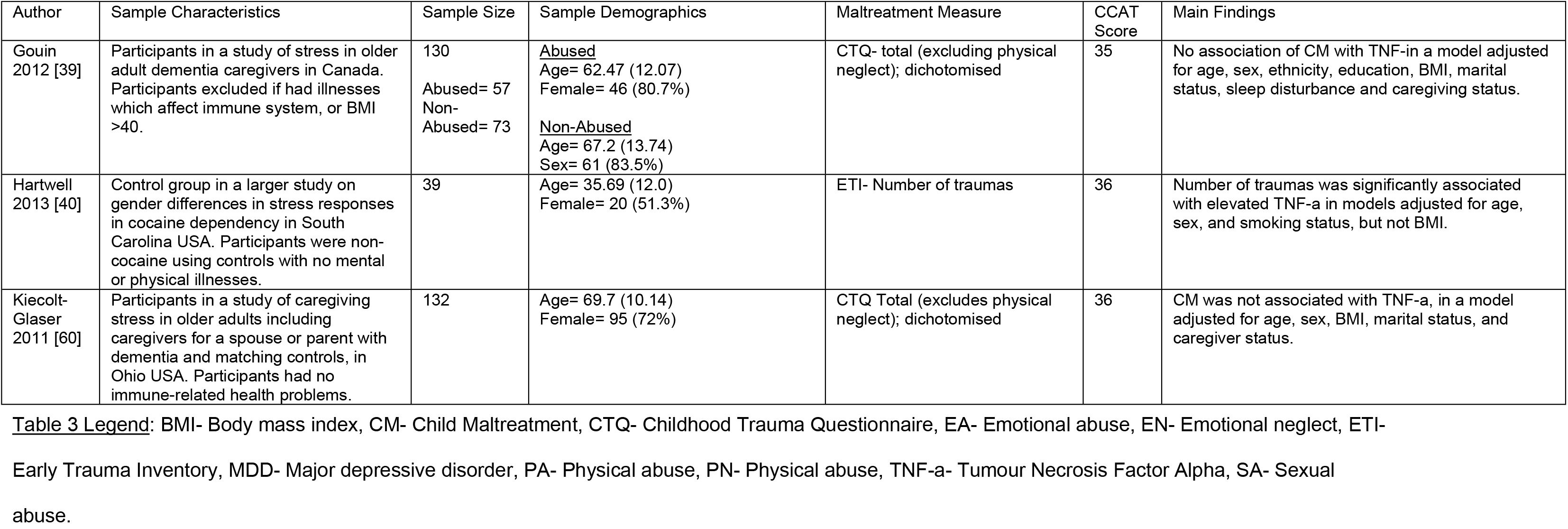
Studies reporting TNF-a: Non-clinical populations

Ten papers, all reporting clinical samples, did not identify a significant association between CM and TNF-a[24–27, 32, 50, 53, 55, 56, 61]. Dennision et al reported elevated levels of TNF-a in participants with schizophrenia and a history of CM compared to participants with schizophrenia only, in an analysis which did not adjust for covariates[54]. Smith et al reported that TNF-a was associated with elevated TNF-a in a sample of 110 African-Americans with and without PTSD, in an analysis which adjusted for age, gender, education, substance use, mental health factors, but not BMI. Toft et al, in a sample of 128 inpatients with MDD, found a significant association between CM (which was not measured using a formal tool) and elevated TNF-a in an unadjusted analysis[62].

In non-clinical samples, two studies did not find a significant association between CM and TNF-a[39, 60]. Hartwell et al in a study of 39 healthy adults in the USA, reported a significant association between the number of traumas on the ETI and elevated TNF-a in an analysis which adjusted for age, sex, and smoking status but not BMI[40].

To summarise most studies did not find a significant association between CM and elevated TNF-a, and none of the studies reporting significant associations had adjusted for BMI.

## Discussion

This systematic review examining the association between CM and markers of systemic inflammation has identified significant variation in the conduct and statistical analysis of studies in this area to the extent that quantitative synthesis of the findings would be invalid. Of note, there was wide variation in how CM exposure was recorded and analysed; for example as a dichotomous versus a continuous variable; as an overall construct versus its subcomponents. Furthermore, the method of analysis varied widely, including between group comparisons, bivariate correlations, linear regression, and more complex modelling. Of note, in analyses where adjustment for covariates was possible there was no consistency in which variables were included. Unsurprisingly, in this context, the findings of studies in this field are inconsistent: the majority of retrospective studies showed no association between CM and inflammatory markers; a number of unadjusted analyses showed statistically significant associations, and a smaller number of fully adjusted analyses showed statistically significant associations but with generally small effect sizes. The variation in conduct and analysis of studies makes it challenging to integrate these disparate findings into a cohesive whole.

This review highlights several limitations in the existing literature. Only four studies (less than 10%) included prospective measures of CM, and these studies only related to CRP. All four of these studies found a significant association between CM and elevated CRP later in life, and two highlighted significant interactions with ethnicity or gender. Baldwin et al have highlighted that retrospective and prospective measures of CM tend to capture different groups of individuals and are not clearly measuring the same construct[63]. This is further supported by the findings of Osborn and colleagues who found that prospective but not retrospective measures of CM were associated with elevated CRP[34]. Despite this small prospective evidence base, and its narrow focus on CRP, the existence of these appropriately adjusted prospective studies demonstrating an association between CM and later increases in CRP suggests that further examination of the links between CM and inflammation is still warranted, but only if studies have sufficient methodological rigour.

In the research base as a whole, studies were inconsistent in their construct of CM: an overall “CM” construct versus sub-types of maltreatment; as a dichotomous variable treating CM as present or absent, or as a continuous measure of the severity of CM. Statistical properties of the way the construct of CM is presented and analysed may contribute to important differences in results (e.g. analyses of continuous measures have more statistical power than dichotomous variables). Studies were also inconsistent in their reporting and analysis of sub-types of abuse. Studies describing results for individual sub-types of CM have reported different effects for different types of maltreatment (most commonly stronger associations of sexual abuse with inflammation)[29, 31, 32, 47, 55]. An approach based on individual sub-types of maltreatment may, however, neglect the inherent complexity and clustering of adversities. For example rather than a specific effect of child sexual abuse as opposed to other maltreatment, the associations found between sexual abuse and inflammation may be more reflective of sexual abuse exposure indexing an overall greater severity of maltreatment exposure and a clustering of multiple adversities[64, 65]. There were limited data on the timing and duration of CM which limits the ability to draw conclusions about sensitive periods in the development of the immune system. Overall, the inconsistencies in measurement of CM could be masking potentially important findings, especially regarding mechanisms.

The conceptualisation and measurement of CM and ACEs more broadly is an area of ongoing debate with relevance to study methodology in this area. Total scores based on the number of forms of CM or ACEs a person has been exposed to is a straight forward way of conceptualising and measuring CM, however it does not reflect the fact that categories of CM or ACE are not equal in their severity or impacts[64]. Analyses based on specific exposures to sub-types of CM or ACE can reflect differential severity and impacts of different types of maltreatment, but fail to reflect the common clustering of maltreatment types (for example intra-familial sexual abuse will almost always be association with physical abuse, emotional abuse, and neglect), and can lose this inherent complexity[64]. Recent work using latent class analysis has identified common clusters of adversity which may represent a better way of conceptualising this area moving forwards[65, 66]. Furthermore it is important to recognise and account for wider forms of adversity which are not fully reflect in traditional conceptions of CM and ACE (which focus more on the immediate family environment), in particular socioeconomic status and wider social adversities such as discrimination[66, 67]. In a similar vein, neurodevelopmental conditions are related to risk of exposure to CM[68] but were not considered as covariates in any of the included studies.

Studies varied greatly in their accounting for potential confounding and mediating variables. Of note, BMI appears to have an important role in the relationship between CM, systemic inflammation and psychopathology. As highlighted previously, most studies finding significant associations between CM and systemic inflammation did not adjust for BMI or related measures (e.g. waist-hip ratio), yet studies employing structural equation modelling suggested that the relationship between CM and inflammation might be mediated by BMI. Based on the current literature it is plausible to speculate that the association between CM and systemic inflammation might be primarily mediated by elevated BMI, but further direct data on this possible association are required.

Most studies were based in Europe or North America and predominantly included participants of white ethnicity. Two studies[35, 46] found significant interactions with ethnicity, contrastingly finding a relationship between neglect and CRP in people of white ethnicity only[35]; and Il-6, fibrinogen, E-Selectin, and sICAM for African-Americans only[46]. Associations between race and health outcomes (particularly in the USA, where there are strong associations between ethnicity and poverty and poor access to healthcare) are likely to be confounded by a range of social and environmental factors[67], and it would be helpful for these apparent associations to be explored more widely and in other settings.

One study[33] found a significant relationship between CM and CRP for females only. Other studies did not directly examine the role of gender, however many studies were conducted exclusively in females or in predominantly female samples which may further affect the overall results. There did not appear to be a consist difference in effects observed in studies limited to participants with psychiatric diagnosis as compared to those limited to community volunteers.

This systematic review is subject to several limitations. Whilst attempts were made to be exhaustive, practical limitations precluded inclusion of foreign language titles and grey literature. Whilst attempting to focus on studies where CM was predominantly child abuse or neglect, limitations in how abuse was measured has meant that some included studies also include broader domains of childhood trauma such as bullying and parental separation. The original protocol for this study aimed to perform a meta-analysis but this was unfortunately neither practicable nor appropriate due to i. significant variation in the exposure concept (CM as a dichotomous or continuous variable; as an overall construct or as sub-components), ii. the use of measurement tools (in particular, difficulties combining between-group comparisons and linear analyses), and iii. inconsistencies in adjustment for covariates where this was done at all. These problems would have significantly impacted the statistical robustness of any findings and potentially created more confusion or, worse, amplified biases, in this already challenging field.

Overall this systematic review has identified an association between CM and elevated CRP in prospective studies, however findings of retrospective studies and for other biomarkers are conflicting. Tentatively at least part of the association between CM and systemic inflammation may be mediated by the association of CM and elevated BMI, which itself may be driven by physiological (such as dysregulated stress-reactivity leading to dysregulation of metabolic pathways) or psychological (such as emotional dysregulation or impulsivity leading to dysregulated eating behaviours) factors, or indeed both. Obesity is strongly associated with low-grade inflammation in a mechanism which may be partially mediated by alterations in the gut microbiome and gut permeability[69], factors which have also been suggested as important drivers of low-grade inflammation and age-related disease[70]. Additional previously unmeasured covariates may also mediate the association of CM with elevated BMI and inflammation, for example neurodevelopmental disorders (which previous work by our group has shown to be associated with obesity[71]) and the gut microbiome, which may mediate the relationship between a range of adverse exposure and inflammation[70]. All of this highlights the importance of applying complex systems methodologies to exploring the interaction of variables holistically and longitudinally[72].

Achieving the research goal of understanding these potentially complex mechanisms would have practical relevance since, if the main mediator is obesity or the gut microbiome, the most effective interventions would likely involve weight loss, exercise, dietary change and early intervention to prevent obesity; whereas if the association between CM and systemic inflammation were more direct, this may point towards a role for anti-inflammatory medications. Further prospective, longitudinal, research using robust and comparable measures of CM with careful consideration of confounding and mediating variables, particularly BMI, are required to bring clarity to this field.

## Supporting Information

S1 File- Details of search strategy

S2 File- PRISMA checklist

S3 File- Systematic review protocol

S1 Table- Details of excluded articles

S2 Table- Results relating to other biomarkers

## References

1. Gilbert R, Widom CS, Browne K, Fergusson D, Webb E, Janson S. Burden and consequences of child maltreatment in high-income countries. The Lancet. 2009;373(9657):68–81.

2. Nemeroff CB. Paradise Lost: The Neurobiological and Clinical Consequences of Child Abuse and Neglect. Neuron. 2016;89(5):892–909. doi: http://dx.doi.org/10.1016/j.neuron.2016.01.019.

3. Hughes K, Bellis MA, Hardcastle KA, Sethi D, Butchart A, Mikton C, et al. The effect of multiple adverse childhood experiences on health: a systematic review and meta-analysis. Lancet Public Health. 2017;2(8):e356–e66. Epub 2017/07/31. doi: 10.1016/S2468-2667(17)30118-4. PubMed PMID: 29253477.

4. Anda RF, Fleisher VI, Felitti VJ, Edwards VJ, Whitfield CL, Dube SR, et al. Childhood Abuse, Household Dysfunction, and Indicators of Impaired Adult Worker Performance. Perm J. 2004;8(1):30–8. doi: 10.7812/tpp/03-089. PubMed PMID: 26704603; PubMed Central PMCID: PMCPMC4690705.

5. Felitti VJ, Anda RF, Nordenberg D, Williamson DF, Spitz AM, Edwards V, et al. Relationship of childhood abuse and household dysfunction to many of the leading causes of death in adults. The Adverse Childhood Experiences (ACE) Study. Am J Prev Med. 1998;14(4):245–58. doi: 10.1016/s0749-3797(98)00017-8. PubMed PMID: 9635069

6. Dube SR, Felitti VJ, Dong M, Giles WH, Anda RF. The impact of adverse childhood experiences on health problems: Evidence from four birth cohorts dating back to 1900. Preventive Medicine: An International Journal Devoted to Practice and Theory. 2003;37(3):268–77. doi: 10.1016/S0091-7435(03)00123-3. PubMed PMID: 2003-07279-010

7. Danese A, Baldwin JR. Hidden Wounds? Inflammatory Links Between Childhood Trauma and Psychopathology. Annual review of psychology. 2017;68:517–44. doi: http://dx.doi.org/10.1146/annurev-psych-010416-044208.

8. Speer K, Upton D, Semple S, McKune A. Systemic low-grade inflammation in post-traumatic stress disorder: a systematic review. J Inflamm Res. 2018;11:111–21. Epub 2018/03/22. doi: 10.2147/JIR.S155903. PubMed PMID: 29606885; PubMed Central PMCID: PMCPMC5868606

9. Danese A, Pariante CM, Caspi A, Taylor A, Poulton R. Childhood maltreatment predicts adult inflammation in a life-course study. Proceedings of the National Academy of Sciences of the United States of America. 2007;104(4):1319–24. doi: http://dx.doi.org/10.1073/pnas.0610362104.

10. Sarwar N, Butterworth AS, Freitag DF, Gregson J, Willeit P, Gorman DN, et al. Interleukin-6 receptor pathways in coronary heart disease: a collaborative meta-analysis of 82 studies. Lancet. 2012;379(9822):1205–13. Epub 2012/03/14. doi: 10.1016/S0140-6736(11)61931-4. PubMed PMID: 22421339; PubMed Central PMCID: PMCPMC3316940

11. Al Bander Z, Nitert MD, Mousa A, Naderpoor N. The Gut Microbiota and Inflammation: An Overview. Int J Environ Res Public Health. 2020;17(20). Epub 2020/10/19. doi: 10.3390/ijerph17207618. PubMed PMID: 33086688; PubMed Central PMCID: PMCPMC7589951

12. Wensley F, Gao P, Burgess S, Kaptoge S, Di Angelantonio E, Shah T, et al. Association between C reactive protein and coronary heart disease: mendelian randomisation analysis based on individual participant data. BMJ. 2011;342:d548. Epub 2011/02/15. doi: 10.1136/bmj.d548. PubMed PMID: 21325005; PubMed Central PMCID: PMCPMC3039696

13. Ridker PM, Libby P, MacFadyen JG, Thuren T, Ballantyne C, Fonseca F, et al. Modulation of the interleukin-6 signalling pathway and incidence rates of atherosclerotic events and all-cause mortality: analyses from the Canakinumab Anti-Inflammatory Thrombosis Outcomes Study (CANTOS). Eur Heart J. 2018;39(38):3499–507. doi: 10.1093/eurheartj/ehy310. PubMed PMID: 30165610

14. Dantzer R, O’Connor JC, Lawson MA, Kelley KW. Inflammation-associated depression: from serotonin to kynurenine. Psychoneuroendocrinology. 2011;36(3):426–36. Epub 2010/10/30. doi: 10.1016/j.psyneuen.2010.09.012. PubMed PMID: 21041030; PubMed Central PMCID: PMCPMC3053088

15. Courtet P, Jaussent I, Genty C, Dupuy AM, Guillaume S, Ducasse D, et al. Increased CRP levels may be a trait marker of suicidal attempt. European Neuropsychopharmacology. 2015;25(10):1824–31. doi: 10.1016/j.euroneuro.2015.05.003. PubMed PMID: 2015-46424-016

16. Ligthart S, Vaez A, Võsa U, Stathopoulou MG, de Vries PS, Prins BP, et al. Genome Analyses of >200,000 Individuals Identify 58 Loci for Chronic Inflammation and Highlight Pathways that Link Inflammation and Complex Disorders. Am J Hum Genet. 2018;103(5):691–706. doi: 10.1016/j.ajhg.2018.09.009. PubMed PMID: 30388399; PubMed Central PMCID: PMCPMC6218410

17. Coelho R, Viola TW, Walss-Bass C, Brietzke E, Grassi-Oliveira R. Childhood maltreatment and inflammatory markers: A systematic review. Acta Psychiatrica Scandinavica. 2014;129(3):180–92. doi: 10.1111/acps.12217. PubMed PMID: 2014-05494-003

18. Baumeister D, Akhtar R, Ciufolini S, Pariante CM, Mondelli V. Childhood trauma and adulthood inflammation: a meta-analysis of peripheral C-reactive protein, interleukin-6 and tumour necrosis factor-alpha. Mol Psychiatry. 2016;21(5):642–9. Epub 2015/06/03. doi: 10.1038/mp.2015.67. PubMed PMID: 26033244; PubMed Central PMCID: PMCPMC4564950

19. Finy MS, Christian LM. Pathways linking childhood abuse history and current socioeconomic status to inflammation during pregnancy. Brain Behav Immun. 2018;74:231–40. Epub 2018/09/16. doi: 10.1016/j.bbi.2018.09.012. PubMed PMID: 30217532; PubMed Central PMCID: PMCPMC6289854

20. Matthews KA, Chang Y-F, Thurston RC, Bromberger JT. Child abuse is related to inflammation in mid-life women: Role of obesity. Brain, Behavior, and Immunity. 2014;36:29–34. doi: 10.1016/j.bbi.2013.09.013. PubMed PMID: 2014-01804-006

21. Moher D, Liberati A, Tetzlaff J, Altman DG, Group P. Preferred reporting items for systematic reviews and meta-analyses: the PRISMA statement. PLoS Med. 2009;6(7):e1000097. Epub 2009/07/21. doi: 10.1371/journal.pmed.1000097. PubMed PMID: 19621072; PubMed Central PMCID: PMCPMC2707599

22. Bernstein DP, Fink L, Handelsman L, Foote J, Lovejoy M, Wenzel K, et al. Initial reliability and validity of a new retrospective measure of child abuse and neglect. Am J Psychiatry. 1994;151(8):1132–6. doi: 10.1176/ajp.151.8.1132. PubMed PMID: 8037246

23. Bremner JD, Vermetten E, Mazure CM. Development and preliminary psychometric properties of an instrument for the measurement of childhood trauma: the Early Trauma Inventory. Depress Anxiety. 2000;12(1):1–12. doi: 10.1002/1520-6394(2000)12:1<1::AID-DA1>3.0.CO;2-W. PubMed PMID: 10999240

24. Counotte J, Bergink V, Pot-Kolder R, Drexhage HA, Hoek HW, Veling W. Inflammatory cytokines and growth factors were not associated with psychosis liability or childhood trauma. PLoS One. 2019;14(7):e0219139. Epub 2019/07/06. doi: 10.1371/journal.pone.0219139. PubMed PMID: 31276524; PubMed Central PMCID: PMCPMC6611659

25. Imai R, Hori H, Itoh M, Lin M, Niwa M, Ino K, et al. Inflammatory markers and their possible effects on cognitive function in women with posttraumatic stress disorder. J Psychiatr Res. 2018;102:192–200. Epub 2018/04/24. doi: 10.1016/j.jpsychires.2018.04.009. PubMed PMID: 29684628

26. Palmos AB, Watson S, Hughes T, Finkelmeyer A, McAllister-Williams RH, Ferrier N, et al. Associations between childhood maltreatment and inflammatory markers. BJPsych Open. 2019;5(1):e3. Epub 2019/02/15. doi: 10.1192/bjo.2018.80. PubMed PMID: 30762500; PubMed Central PMCID: PMCPMC6343120

27. Zeugmann S, Quante A, Popova-Zeugmann L, Kössler W, Heuser I, Anghelescu I. Pathways linking early life stress, metabolic syndrome, and the inflammatory marker fibrinogen in depressed inpatients. Psychiatria Danubina. 2012;24(1):57–65. PubMed PMID: 2012-09083-007

28. de Punder K, Entringer S, Heim C, Deuter CE, Otte C, Wingenfeld K, et al. Inflammatory Measures in Depressed Patients With and Without a History of Adverse Childhood Experiences. Front Psychiatry. 2018;9:610. Epub 2018/12/13. doi: 10.3389/fpsyt.2018.00610. PubMed PMID: 30538644; PubMed Central PMCID: PMCPMC6277546

29. Aas M, Dieset I, Hope S, Hoseth E, Morch R, Reponen E, et al. Childhood maltreatment severity is associated with elevated C-reactive protein and body mass index in adults with schizophrenia and bipolar diagnoses. Brain Behav Immun. 2017;65:342–9. Epub 2017/06/18. doi: 10.1016/j.bbi.2017.06.005. PubMed PMID: 28619247

30. Fanning JR, Lee R, Gozal D, Coussons-Read M, Coccaro EF. Childhood trauma and parental style: Relationship with markers of inflammation, oxidative stress, and aggression in healthy and personality disordered subjects. Biological Psychology. 2015;112:56–65. doi: 10.1016/j.biopsycho.2015.09.003. PubMed PMID: 2015-52970-009

31. Hepgul N, Pariante CM, Dipasquale S, DiForti M, Taylor H, Marques TR, et al. Childhood maltreatment is associated with increased body mass index and increased C-reactive protein levels in first-episode psychosis patients. Psychol Med. 2012;42(9):1893–901. Epub 2012/01/21. doi: 10.1017/s0033291711002947. PubMed PMID: 22260948; PubMed Central PMCID: PMCPMC4081598

32. Quide Y, Bortolasci CC, Spolding B, Kidnapillai S, Watkeys OJ, Cohen-Woods S, et al. Association between childhood trauma exposure and pro-inflammatory cytokines in schizophrenia and bipolar-I disorder. Psychol Med. 2019;49(16):2736–44. Epub 2018/12/19. doi: 10.1017/s0033291718003690. PubMed PMID: 30560764

33. Baldwin JR, Arseneault L, Caspi A, Fisher HL, Moffitt TE, Odgers CL, et al. Childhood victimization and inflammation in young adulthood: A genetically sensitive cohort study. Brain, behavior, and immunity. 2018;67:211–7. doi: https://dx.doi.org/10.1016/j.bbi.2017.08.025.

34. Osborn M, Widom CS. Do documented records and retrospective reports of childhood maltreatment similarly predict chronic inflammation? Psychol Med. 2019:1–10. Epub 2019/09/24. doi: 10.1017/s0033291719002575. PubMed PMID: 31544727

35. Nikulina V, Widom CS. Do race, neglect, and childhood poverty predict physical health in adulthood? A multilevel prospective analysis. Child Abuse & Neglect. 2014;38(3):414–24. doi: 10.1016/j.chiabu.2013.09.007. PubMed PMID: 2013-38953-001

36. Anderson EL, Fraser A, Caleyachetty R, Hardy R, Lawlor DA, Howe LD. Associations of adversity in childhood and risk factors for cardiovascular disease in mid-adulthood. Child Abuse & Neglect. 2018;76:138–48. doi: 10.1016/j.chiabu.2017.10.015. PubMed PMID: 2018-04158-015

37. Carpenter LL, Gawuga CE, Tyrka AR, Price LH. C-reactive protein, early life stress, and wellbeing in healthy adults. Acta Psychiatr Scand. 2012;126(6):402–10. Epub 2012/06/12. doi: 10.1111/j.1600-0447.2012.01892.x. PubMed PMID: 22681496; PubMed Central PMCID: PMCPMC3580169

38. Gong J, Wang Y, Liu J, Fu X, Cheung EFC, Chan RCK. The interaction between positive schizotypy and high sensitivity C-reactive protein on response inhibition in female individuals. Psychiatry Res. 2019;274:365–71. Epub 2019/03/11. doi: 10.1016/j.psychres.2019.02.064. PubMed PMID: 30852429

39. Gouin J-P, Glaser R, Malarkey WB, Beversdorf D, Kiecolt-Glaser JK. Childhood abuse and inflammatory responses to daily stressors. Annals of Behavioral Medicine. 2012;44(2):287–92. doi: 10.1007/s12160-012-9386-1. PubMed PMID: 2012-25446-016

40. Hartwell KJ, Moran-Santa Maria MM, Twal WO, Shaftman S, DeSantis SM, McRae-Clark AL, et al. Association of elevated cytokines with childhood adversity in a sample of healthy adults. Journal of Psychiatric Research. 2013;47(5):604–10. doi: 10.1016/j.jpsychires.2013.01.008. PubMed PMID: 2013-09208-004

41. Kim S, Watt T, Ceballos N, Sharma S. Adverse childhood experiences and neuroinflammatory biomarkers-The role of sex. Stress Health. 2019;35(4):432–40. Epub 2019/05/18. doi: 10.1002/smi.2871. PubMed PMID: 31099473

42. Powers A, Michopoulos V, Conneely K, Gluck R, Dixon H, Wilson J, et al. Emotion Dysregulation and Inflammation in African-American Women with Type 2 Diabetes. Neural Plasticity. 2016;2016:8926840. doi: http://dx.doi.org/10.1155/2016/8926840.

43. Boeck C, Koenig AM, Schury K, Geiger ML, Karabatsiakis A, Wilker S, et al. Inflammation in adult women with a history of child maltreatment: The involvement of mitochondrial alterations and oxidative stress. Mitochondrion. 2016;30:197–207. Epub 2016/08/18. doi: 10.1016/j.mito.2016.08.006. PubMed PMID: 27530300

44. Thurston RC, Chang Y, Barinas-Mitchell E, Von Kanel R, Richard Jennings J, Santoro N, et al. Child abuse and neglect and subclinical cardiovascular disease among midlife women. Psychosomatic Medicine. 2017;79(4):441–9. doi: http://dx.doi.org/10.1097/PSY.0000000000000400.

45. Pinto Pereira SM, Stein Merkin S, Seeman T, Power C. Understanding associations of early-life adversities with mid-life inflammatory profiles: Evidence from the UK and USA. Brain Behav Immun. 2019;78:143–52. Epub 2019/01/27. doi: 10.1016/j.bbi.2019.01.016. PubMed PMID: 30682500; PubMed Central PMCID: PMCPMC6941353

46. Slopen N, Lewis TT, Gruenewald TL, Mujahid MS, Ryff CD, Albert MA, et al. Early life adversity and inflammation in African Americans and Whites in the Midlife in the United States Survey. Psychosomatic Medicine. 2010;72(7):694–701. doi: 10.1097/PSY.0b013e3181e9c16f. PubMed PMID: 2011-26515-012

47. Bertone-Johnson ER, Whitcomb BW, Missmer SA, Karlson EW, Rich-Edwards JW. Inflammation and early-life abuse in women. American Journal of Preventive Medicine. 2012;43(6):611–20. doi: 10.1016/j.amepre.2012.08.014. PubMed PMID: 2012-31219-008

48. Rooks C, Veledar E, Goldberg J, Bremner JD, Vaccarino V. Early trauma and inflammation: Role of familial factors in a study of twins. Psychosomatic Medicine. 2012;74(2):146–52. doi: http://dx.doi.org/10.1097/PSY.Ob013e318240a7d8.

49. Schrepf A, Markon K, Lutgendorf SK. From childhood trauma to elevated c-reactive protein in adulthood: The role of anxiety and emotional eating. Psychosomatic Medicine. 2014;76(5):327–36. doi: http://dx.doi.org/10.1097/PSY.0000000000000072.

50. Corsi-Zuelli F, Loureiro CM, Shuhama R, Fachim HA, Menezes PR, Louzada-Junior P, et al. Cytokine profile in first-episode psychosis, unaffected siblings and community-based controls: the effects of familial liability and childhood maltreatment. Psychol Med. 2019:1–9. Epub 2019/05/09. doi: 10.1017/s0033291719001016. PubMed PMID: 31064423

51. Porcu M, Machado RCBR, Urbano M, Verri WA, Rossaneis AC, Vargas HO, et al. Depressed female smokers have higher levels of soluble tumor necrosis factor receptor 1. Addictive Behaviors Reports. 2018;7:90–5. doi: http://dx.doi.org/10.1016/j.abrep.2018.03.004.

52. Smith AK, Conneely KN, Kilaru V, Mercer KB, Weiss TE, Bradley B, et al. Differential immune system DNA methylation and cytokine regulation in post-traumatic stress disorder. American Journal of Medical Genetics Part B: Neuropsychiatric Genetics. 2011;156(6):700–8. doi: 10.1002/ajmg.b.31212. PubMed PMID: 2014-15537-008

53. Lu S, Peng H, Wang L, Vasish S, Zhang Y, Gao W, et al. Elevated specific peripheral cytokines found in major depressive disorder patients with childhood trauma exposure: A cytokine antibody array analysis. Comprehensive Psychiatry. 2013;54(7):953–61. doi: 10.1016/j.comppsych.2013.03.026. PubMed PMID: 2013-15209-001

54. Dennison U, McKernan D, Cryan J, Dinan T. Schizophrenia patients with a history of childhood trauma have a pro-inflammatory phenotype. Psychol Med. 2012;42(9):1865–71. Epub 2012/02/24. doi: 10.1017/s0033291712000074. PubMed PMID: 22357348

55. Grosse L, Ambrée O, Jörgens S, Jawahar MC, Singhal G, Stacey D, et al. Cytokine levels in major depression are related to childhood trauma but not to recent stressors. Psychoneuroendocrinology. 2016;73:24–31. doi: 10.1016/j.psyneuen.2016.07.205. PubMed PMID: 2016-46963-005

56. Pedrotti Moreira F, Wiener CD, Jansen K, Portela LV, Lara DR, Souza LDDM, et al. Childhood trauma and increased peripheral cytokines in young adults with major depressive: Population-based study. Journal of Neuroimmunology. 2018;319:112–6. doi: http://dx.doi.org/10.1016/j.jneuroim.2018.02.018.

57. Muller N, Krause D, Barth R, Myint AM, Weidinger E, Stettinger W, et al. Childhood Adversity and Current Stress are related to Pro-and Anti-inflammatory Cytokines in Major Depression. J Affect Disord. 2019;253:270–6. Epub 2019/05/08. doi: 10.1016/j.jad.2019.04.088. PubMed PMID: 31063941

58. Munjiza A, Kostic M, Pesic D, Gajic M, Markovic I, Tosevski DL. Higher concentration of interleukin 6-A possible link between major depressive disorder and childhood abuse. Psychiatry Research. 2018;264:26–30. doi: http://dx.doi.org/10.1016/j.psychres.2018.03.072.

59. Davis MC, Lemery-Chalfant K, Yeung EW, Luecken LJ, Zautra AJ, Irwin MR. Interleukin-6 and Depressive Mood Symptoms: Mediators of the Association Between Childhood Abuse and Cognitive Performance in Middle-Aged Adults. Ann Behav Med. 2019;53(1):29–38. Epub 2018/03/22. doi: 10.1093/abm/kay014. PubMed PMID: 29562248; PubMed Central PMCID: PMCPMC6301312

60. Kiecolt-Glaser JK, Gouin J-P, Weng N-P, Malarkey WB, Beversdorf DQ, Glaser R. Childhood adversity heightens the impact of later-life caregiving stress on telomere length and inflammation. Psychosomatic Medicine. 2011;73(1):16–22. doi: 10.1097/PSY.0b013e31820573b6. PubMed PMID: 2011-26702-003

61. Kraav SL, Tolmunen T, Karkkainen O, Ruusunen A, Viinamaki H, Mantyselka P, et al. Decreased serum total cholesterol is associated with a history of childhood physical violence in depressed outpatients. Psychiatry Res. 2019;272:326–33. Epub 2019/01/01. doi: 10.1016/j.psychres.2018.12.108. PubMed PMID: 30597385

62. Toft H, Neupane SP, Bramness JG, Tilden T, Wampold BE, Lien L. The effect of trauma and alcohol on the relationship between level of cytokines and depression among patients entering psychiatric treatment. BMC Psychiatry. 2018;18(1):95. Epub 2018/04/11. doi: 10.1186/s12888-018-1677-z. PubMed PMID: 29631540; PubMed Central PMCID: PMCPMC5891976

63. Baldwin JR, Reuben A, Newbury JB, Danese A. Agreement Between Prospective and Retrospective Measures of Childhood Maltreatment: A Systematic Review and Meta-analysis. JAMA Psychiatry. 2019;76(6):584–93. doi: 10.1001/jamapsychiatry.2019.0097. PubMed PMID: 30892562; PubMed Central PMCID: PMCPMC6551848

64. Lacey RE, Minnis H. Practitioner Review: Twenty years of research with adverse childhood experience scores-Advantages, disadvantages and applications to practice. J Child Psychol Psychiatry. 2020;61(2):116–30. Epub 2019/10/14. doi: 10.1111/jcpp.13135. PubMed PMID: 31609471

65. Lacey RE, Pinto Pereira SM, Li L, Danese A. Adverse childhood experiences and adult inflammation: Single adversity, cumulative risk and latent class approaches. Brain Behav Immun. 2020. Epub 2020/03/24. doi: 10.1016/j.bbi.2020.03.017. PubMed PMID: 32201253

66. Lacey RE, Howe LD, Kelly-Irving M, Bartley M, Kelly Y. The Clustering of Adverse Childhood Experiences in the Avon Longitudinal Study of Parents and Children: Are Gender and Poverty Important? J Interpers Violence. 2020:886260520935096. Epub 2020/07/08. doi: 10.1177/0886260520935096. PubMed PMID: 32639853

67. Williams DR, Priest N, Anderson NB. Understanding associations among race, socioeconomic status, and health: Patterns and prospects. Health Psychol. 2016;35(4):407–11. doi: 10.1037/hea0000242. PubMed PMID: 27018733; PubMed Central PMCID: PMCPMC4817358

68. Dinkler L, Lundström S, Gajwani R, Lichtenstein P, Gillberg C, Minnis H. Maltreatment-associated neurodevelopmental disorders: a co-twin control analysis. J Child Psychol Psychiatry. 2017;58(6):691–701. Epub 2017/01/17. doi: 10.1111/jcpp.12682. PubMed PMID: 28094432

69. Cox AJ, West NP, Cripps AW. Obesity, inflammation, and the gut microbiota. Lancet Diabetes Endocrinol. 2015;3(3):207–15. Epub 2014/07/22. doi: 10.1016/S2213-8587(14)70134-2. PubMed PMID: 25066177

70. Shiels PG, Buchanan S, Selman C, Stenvinkel P. Allostatic load and ageing: linking the microbiome and nutrition with age-related health. Biochem Soc Trans. 2019;47(4):1165–72. Epub 2019/08/15. doi: 10.1042/BST20190110. PubMed PMID: 31416886

71. Mina TH, Lahti M, Drake AJ, Räikkönen K, Minnis H, Denison FC, et al. Prenatal exposure to very severe maternal obesity is associated with adverse neuropsychiatric outcomes in children. Psychol Med. 2017;47(2):353–62. Epub 2016/10/25. doi: 10.1017/S0033291716002452. PubMed PMID: 27776561

72. Ioannidis K, Askelund AD, Kievit RA, van Harmelen AL. The complex neurobiology of resilient functioning after childhood maltreatment. BMC Med. 2020;18(1):32. Epub 2020/02/14. doi: 10.1186/s12916-020-1490-7. PubMed PMID: 32050974; PubMed Central PMCID: PMCPMC7017563

